# Improving the reliability of molecular string representations for generative chemistry

**DOI:** 10.1101/2024.10.07.617002

**Authors:** Etienne Reboul, Zoe Wefers, Harish Prabakaran, Jérôme Waldispühl, Antoine Taly

## Abstract

Generative modeling for chemistry has advanced rapidly in recent years, but this surge in popularity raises a foundational question: which molecular representation is best suited for modern machine learning models? Despite not being designed for generative tasks, SMILES remain the most commonly used string-based representation. However, while SMILES follow strict syntactic rules, grammatically correct SMILES strings do not always correspond to valid molecules. SELFIES were introduced as an alternative that addresses this limitation by ensuring that every string of SELFIES tokens represents to a valid molecule.

In this study, we comprehensively evaluate the limitations of both SMILES and SELFIES as representations for generative models. We define two key criteria for robust molecular generation: viability, generated strings represent novel, unique molecules with correct valence, and fidelity, the distribution of physicochemical properties from sampled molecules resembles that of the training data. We find that approximately one-fifth of molecules generated using canonical SMILES are invalid, failing the viability criterion. In contrast, all SELFIES-generated molecules are viable, but they deviate significantly from the training distribution, indicating low fidelity.

To address these limitations, we develop data augmentation procedures for both representations. While simplifying the SELFIES grammar yields only modest gains in fidelity, our stochastic augmentation method for SMILES, ClearSMILES, significantly improves both viability and fidelity. ClearSMILES simplifies syntax by reducing the vocabulary size and explicitly encoding aromaticity via Kekulé SMILES, making it easier string representations for models to process. Using ClearSMILES, the rate of invalid samples decreases by an order of magnitude, from 20% to 2.2%, and fidelity to the training distribution is also moderately improved.

Generative chemistry has seen rapid development recently. However, models based on string representations of molecules still rely largely on SMILES^1^ that have not been developed for this context and SELFIES ^2^ who were introduced to reduce those problems. The goal of this study is to first analyze the difficulty encountered by a small generative model when using SMILES and SELFIES.

Our study found that SELFIES and canonical SMILES ^3^ are not fully reliable representations for a small generative model, i.e. do not ensure concurrently the viability and fidelity of samples. Viable samples represent novel, unique molecules with correct valence, while fidelity is efficient distribution learning of key physico-chemical properties. ^4^ In fact, 20% of the samples generated using canonical SMILES input representation do not correspond to valid molecules. In contrast, samples generated using SELFIES were all viable but where not able to reproduce as well the distribution of physico-chemical properties as SMILES.

As a mitigation strategy for the previously identified problems, we have developed data augmentation procedures for both SELFIES and SMILES. Simplifying the complex syntax of SELFIES yielded only marginal improvements in string stability and overall fidelity to the training set. For SMILES, we developed a stochastic data augmentation procedure called ClearSMILES, which reduces the vocabulary size needed to represent a SMILES dataset, explicitly represents aromaticity via Kekulé SMILES, ^3^ and reduces the effort required by deep learning models to process SMILES. ClearSMILES reduced the rate of invalid samples by an order of magnitude, from 20% to 2.2%, and improved the fidelity of samples to the training set.

## 1 Introduction

Traditional *In silico* techniques for de novo drug design are based on virtual screening (VS) of large virtual libraries. The hits found by the VS are purchased or synthesized in order to experimentally confirm their activity. The validated compounds are then optimized by specialists of various field, including medicinal chemists, toxicologists or data scientists, to achieve a desired profile of biological properties, that is, pharmacokinetics, toxicity and activity. The traditional pipeline is as time-consuming as it is expensive. The goal of generative chemistry is to use machine learning (ML) to generate new compounds by sampling from a known distribution (unconditional sampling), possibly by constraining the sampling with desired properties (conditional sampling) to match the desired chemical profile.^5^ A popular class of generative models is the variational autoencoder (VAE).^2,6–9^

There are two predominant molecular representations for a VAE: graphs and string. ^10^ Molecules are most naturally represented by 2D graph data structures, as they can be assimilated to a graph where atoms are vertices and chemical bonds are edges. Therefore, a 2D graph representation explicitly provides structural information through separate adjacency matrices and one-hot encoded matrices for atom and bond attributes. This explicit separation of structural elements (connectivity, atom types, bond types) provides directly accessible information to the model. However, preprocessing and postprocessing 2D graphs can be slow, making parsing of a large batch of molecules using python packages such as RDKit^11^ quite cumbersome compared to their string counterpart.

String representations are linear and easy to manipulate. They can be made model-readable by converting the sequence of tokens to a unique one-hot encoding matrix. A one-hot encoded matrix of a string representation is more memory friendly but requires the model to implicitly extract the molecular structure. The early models using string representation performed poorly compared to the graph-based model counterpart. ^12^ The performance gap was bridged by piggybacking on advances in natural language processing (NLP), especially by using a transformer-like model.^7,13,14^ Currently, the state-of-the-art for a string-based generative model performs roughly as well as the graph-based one. ^15^

The last decades have seen the emergence of multiple string molecular representations dedicated to chemo-informatics usage. The oldest and most established string representation of molecules is SMILES.^1^ It is widely used to store chemical structures in virtual online databases and as input to a multitude of chemo-informatics tools, including the staples of the open-source community such as RDKIT^16^ and openbabel. ^17^ SMILES syntax is very simple and human-readable (Figure 1). Atoms are represented by their corresponding atomic symbols in the periodic table. They form the backbone of the molecule from which branches sprout to represent nonlinear chemical patterns, such as ester groups. They are represented by closed matching parentheses. Matching pairs of digits placed after the atom tokens are used to represent the connection in the molecular graph of atoms that are nonadjacent in the string, thereby forming rings.

**Figure 1:**
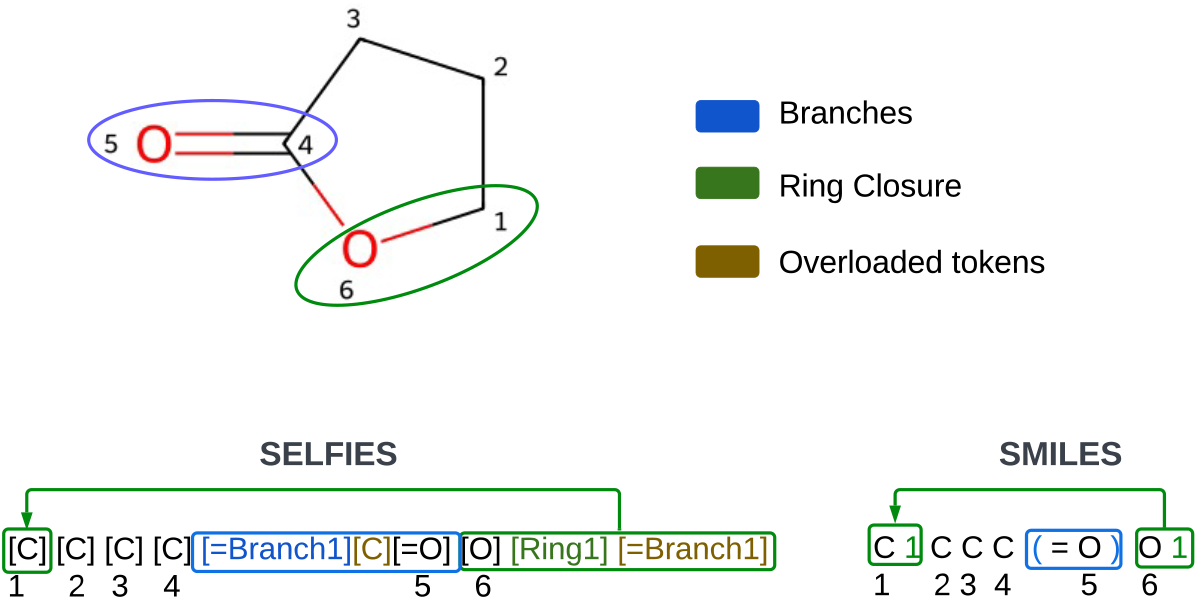
Example of SMILES and SELFIES for gamma-butyroLactone (GBL). The branch of butyrolactone is cirled in blue on the GBL’s 2D structure and boxed in blue in the SELFIES and SMILES examples. The ring closure is circled in green on GBL’s 2D structure and boxed in green also in the SELFIES and SMILES examples. The number displayed in the figure represent the arbitrary indexing of GBL’s atoms.

Although SMILES are the most commonly used string representation of molecules, they are not robust. Every string of characters that complies with the SMILES branch and ring grammar does not necessarily represent a valid molecule. This usually happens when one or more of the bonds implied by the SMILES string break the valence of one of the atoms. SELFIES^2,18^ were created on top of SMILES such that every string of SELFIES tokens can be translated into SMILES that represent a valid molecule. This was achieved via a rigorous and fully defined algorithm, which deletes erroneous ring, branch, and atom tokens that would imply a break of the valence of any atom. For this reason, it is stated that every string of SELFIES tokens represents a valid molecule. This is particularly advantageous in the context of machine learning, as it guarantees that the output has a valid valence. However, it does not guarantee that the generated molecules will have properties similar to those of the input molecules, such as druglikeness. The SELFIES grammar diverges significantly from the SMILES grammar. Unlike SMILES, SELFIES do not utilize pairs of digits or parenthesis tokens to represent rings and branches (Figure 1). Instead, they employ branch and ring tokens to indicate the positions of branches and rings. The sizes of branches and rings are specified by a number of overloaded tokens added after the branch or ring token. In this context, overloaded tokens are interpreted as numbers instead of their original meaning; the map to a numerical value is shown in the Supplementary Table 1.

There is a current trend in NLP for models to become bigger and more expensive to train.^19^ We have chosen instead a simple RNNAttn-VAE^7^ as our generative model for the following reason: it is faster and cheaper to train due to its simple architecture; the single-head attention used in the decoder will be simpler to interpret, and its simplicity will make it easier to pinpoint the difficulty the neural network has in grasping the grammar of SELFIES and SMILES. We also make the assumption that a simple model such as the RNNAttn-VAE is less likely to overfit the data, with better generalization and reduce the risk of simple memorization. The short-term aim is to have a small generative model like RNN based VAE with narrower performance gap compared to more complex models that are transformer-based. In the long term, any larger model using the progresses proposed below should only benefit from them.

The main avenue we are planning to improve is the reliability of string molecular representation. Reliability for generative chemistry can be understood as the combination of viability and fidelity. A reliable string representation enables a generative model to generate novel and unique compounds with valid valence. Furthermore, a reliable string representation allows a generative model to learn the distribution of physico-chemical properties of the training sets.

For that purpose, this study is centered around two parts: The first part is to assess what goes wrong with the generation of samples using SELFIES and SMILES as the representation for our RNNAttn-VAE. The second part is centered around mitigating strategies by developing data augmentation procedures for both SMILES and SELFIES.

## 2 Methods

### 2.1 Datasets

The different models were trained using the MOSES database.^12^ The MOSES database is a subset of the ZINC15^20^ Clean Leads collection that was designed to be representative of drug-like compounds. To that effect, the selected molecules were filtered using the following criteria: a weight between 250 to 350 Da, the number of rotatable bonds equal to or below seven, no charged atoms in the molecule (e.g. no nitrate group), only molecules with C, N, S, O, F, Cl, Br, and H atoms were kept, and no molecules with cycles longer than eight atoms. The MOSES database consists of a training set (1.7M molecules), test set (176K molecules), and scaffold test set (176K molecules) of molecules with unique scaffolds that never appear in the training set.

### 2.2 RNNAttn-VAE

A VAE is made up of two parts: an encoder and a decoder.^21^ The encoder takes an input array *x* and compresses it to the latent space of *d*_*latent*_ dimensions (where *d*_*latent*_*<d*_*input*_). Specifically, the encoder outputs two parameters: *µ* represents the mean, *σ* represents the standard deviation. They are used to produce the array *z. z* is defined with the equation 1. Where and *ϵ* is a random value between 0 and 1.

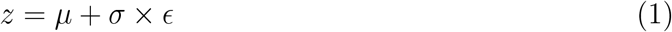

The sample *z* represents compressed information that contains key features of the input *x* and is used as input for the decoder to reconstruct *x*^*′*^, as close as can be to a copy of *x*. By making the mean and variance of the latent space follow a known distribution, the trained decoder can be used to generate new samples from random noise generated from the known distribution.

Our code was developed from that of RNNAttn-VAE.^7^ The original base code is available on github: https://github.com/oriondollar/TransVAE. Briefly, our encoder and decoder are composed of 126 Gated Recurrent Unit (GRU) layer with Batch Norm repeated 3 times. A single attention head is placed after the last GRU layer of the encoder, followed by a convolutional bottleneck. The latent space of our VAE is composed of 15 or 22 latent dimensions. A Softmax function is used to obtain the output of the model from the output of the decoder.

SMILES and SELFIES were tokenized and then converted to a one-hot encoded matrix of dimensions *n × m* where *n* is the number of tokens in the longest SMILES/SELFIES and *m* is the number of possible tokens. This was used as input, and each model was trained during 100 epochs.

The RNNAttn-VAE has a loss function that combines two losses: the reconstruction loss and the Kullback-Leibler divergence (KLD). A weighted cross-entropy loss is used as the reconstruction loss to measure how well the generated output matches the input data. The weights are inversely proportional to the occurrence. The exact scheme to compute the cross-entropy weights can be found at this location on github. Meanwhile, the KLD acts as a regularizer, ensuring that the learned latent space approximates a Gaussian distribution. It measures the divergence between the approximate posterior distribution and the prior distribution, guiding the model to learn a smooth and continuous latent space representation.

### 2.3 Metrics

#### 2.3.1 fidelity metrics

The fidelity assessment, i.e. accurate distribution learning,^4^ of the compound was based on the following metrics: quantitative estimate of drug-likeness (QED), ^22^ synthetic accessibility estimation (SA), ^23^ molecular weight (MW), topological polar surface area (TPSA^24^). All metrics were computed using the RDKIT toolkit (version 2023.09.1).

#### 2.3.2 samples uniqueness and novelty

The uniqueness for valid samples was calculated using the IUPAC international chemical identifier (InChI).^25^ This ensures that molecules generated from samples are actually unique, as a given molecule can have a myriad of valid SMILES. The novelty rate was computed by computing the size of the intersection between each InChI’s of samples and InChI’s of training set.

#### 2.3.3 SMILES validity

The validity, specifically the proper valence (semantic validity) and the respect of SMILES grammar (syntactical validity), of the molecules generated using SMILES was tested using RDKit. A molecule is considered valid if RDKIT can generate a 2D structure from the generated SMILES. Validity rates are calculated as the ratio of invalid molecules to total molecules and expressed as a percentage. For invalid molecules, the cause of invalidity is determined by parsing the error message and classifying it into the following categories that are similar to those found in the literature: ^10,26^ aromaticity error, ring error, parenthesis error, valence error and syntax error. Ring errors, parenthesis errors, aromaticity errors, and syntax errors are syntactic errors, whereas valence errors are semantic errors.

#### 2.3.4 SELFIES string stability

The decoding algorithm SELFIES will invariably output a valid SMILES, with correct valence. Therefore, we developed a proxy validity metric called string stability, as a reflection of the model’s ability to learn and reproduce the SELFIES grammar. A SELFIES is considered stable if the regenerated SELFIES obtained by decoding the original SELFIES to SMILES and then re-encoding it back to SELFIES is identical to the original SELFIES, i.e. a lossless encoding-decoding. This is achieved using the selfies Python module (version 2.1). If a sequence of SELFIES tokens has an incorrect valence it will be detected by this metric.

For instance, Let’s take a SELFIES of dichloromethane: [Cl][C][C][Cl], if we mutate it into [Cl][Cl][C][Cl] the mutated tokens sequence has a wrong valence. The mutated SELFIES implies that the newly introduced chlorine atom is bounded to both the previous chlorine atom and the next carbon atom. When the mutatedSELFIES is decoded to SMILES, the SELFIES decoding algorithm truncates the sequence of atoms to ClCl or chlorine gas.

To ascertain that the SELFIES encoding algorithm doesn’t make arbitrary changes in the SELFIES token sequence, we made a negative control with the MOSES dataset. The protocol is as follows:

1. Download the MOSES dataset
2. Encode all SMILES to SELFIES
3. test the stability with 10 replica per SELFIES

The code is available in a Jupyter Notebook distributed with Google Colab.

To further quantify the loss of information related to the SELFIES instability, we introduced a proxy loss based on the amount of token lost between the original and regenerated SELFIES. The token loss is the weighted difference between the number of tokens in the regenerated SELFIES (*n*_regenerated_) and the original SELFIES (*n*_original_), as defined in equation 2:

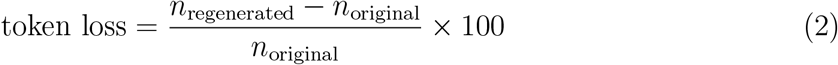

To identify the root cause of the SELFIES instability, we developed a pipeline based on the alignment of original and regenerated SELFIES to highlight the transformation made by the SELFIES algorithm.

First, a regenerated SELFIES is obtained for the string stability test. Both the original SELFIES and regenerated SELFIES are tokenized to be labeled in 4 different categories: a for the atom tokens, b for the branch tokens, r for the ring tokens, and n for the over-loaded/numerical tokens.

The labels and the tokens are then merged to get annotated tokens. Those annotated tokens are then aligned using the Needleman Wunsch algorithm implemented in the string2string (0.0.150) Python module. The mutations detected by alignment were classified as the 3 standard mutations in biology: insertion, deletion, and substitution.

Note that the alignments are not perfect. When decoding, the SELFIES algorithm can remove, delete, or change the length of the ring and branches. The Needleman-Wunsch algorithm will detect the modification but will map the token incorrectly when substitution and addition happen simultaneously.

#### 2.3.5 Shannon Information Entropy

The Shannon information entropy defined in equation 3 is used to measure the dispersion of information in the latent space.

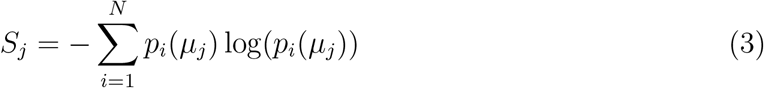

### 2.4 Data augmentation

#### 2.4.1 SMILES randomization parameter selection

For the first 15,252 SMILES in the MOSES dataset, we have applied the following protocol to model the relationship between the number of randomized smiles and the number of unique randomized smiles: We split the computation of 1 million randomized SMILES for each molecule into chunks of 1000 randomized SMILES. A set records the cumulative unique smiles yield by each chunk. If the size of the set of unique smiles does not increase after 10 consecutive chunks, the calculation is prematurely stopped.

#### 2.4.2 ClearSMILES

ClearSMILES is a stochastic data augmentation procedure that aims to simultaneously achieve dimensionality reduction and minimize the global attention effort required for processing a SMILES. ClearSMILES leverages two properties of SMILES: the level of the SMILES representation and the uniqueness problem of SMILES.

The traditional representation of aromatic atoms in SMILES uses lowercase symbols. This representation abstracts away the choice of a particular mesomer and faithfully represents the concept of electron delocalization in the aromatic system. It has a main drawback: the introduction of a new set of tokens: ‘[nH]’,’c’,’n’, ‘o’, ‘s’. The relationship between aliphatic and non-aliphatic tokens has to be learned or inferred, which is most likely not a trivial task. The kekule representation^3^ picks an arbitrary mesomer which enables the representation of aromatic cycles with aliphatic symbols and the electronic system as patterns of double bonds. Dropping the aromatic token from the vocabulary will lead to a reduction in the dimensionality of the one-hot encoded matrix with a smaller vocabulary, although SMILES are longer due to double bond tokens introduced by the Kekule representation.

The second aspect that is tackled by ClearSMILES is the uniqueness problem. A plethora of SMILES can represent the same molecule because each path through the 2D graph of a molecule will yield a different SMILES. The long range dependencies in SMILES, i.e., digit pairs and branches matching parenthesis, can vary in size and in number (branches only) depending on the SMILES. Therefore one of the key mechanism behind ClearSMILES is to find a SMILES with limited long range dependencies.

The ClearSMILES pipeline can be divided into two main parts: generation and filtration. The first part consists of generating 100,000 randomized Kekule SMILES. The sampling of a large number of SMILES is used to ensure that the large space of possible SMILES per molecule is sampled exhaustively. The second part of the pipeline is the sequential filtering of randomized Kekule SMILES generated in the first part. The filtration protocol is as follow:

1. Removing duplicates of randomized Kekule SMILES
2. Selecting SMILES with the lowest maximum digits
3. Selecting SMILES with the lowest memory score
4. Selecting the first SMILES string after sorting SMILES alphanumerically

The first filtration step is to prevent the waste of computational resources on redundant randomized Kekule SMILES. For the second step of the filtration part, we filter the unique SMILES to keep only those with the lowest maximum digit. This serves a dual purpose. It limits the number of digit tokens needed to describe the rings in SMILES, achieving secondary dimensionality reduction of the one hot-encoding. It also guarantees that the selected randomized SMILES exhibit maximum disentanglement between rings, as depicted in **Figure 3 (b)**).

In the fourth step of the filtration part, we calculate the semantic memory map for each remaining SMILES. The semantic memory map generation process tracks the number of open semantic features, i.e. branches and ring closures, at each position in a SMILES string. To build this map, we first tokenize the SMILES string and initialize an array of zeros matching the number of tokens. We then iterate through each token, keeping track of branch and ring openings/closings. When we encounter an opening parenthesis ‘(‘, we increment the value at the current position by 1. For a closing parenthesis ‘)’, we decrement by 1. Digits representing ring closures are handled using a set: when a digit appears for the first time, it represents a ring opening (increment by 1); when the same digit appears again, it represents a ring closing (decrement by 1). Finally, we compute and return the cumulative sum of this array, which effectively represents the number of open semantic features after processing each token. The pseudo code for this procedure is described in algorithm 1.

The Memory score of the SMILES is then calculated as the arithmetic mean of the memory map. The SMILES with the minimum memory score are ClearSMILES. The objective of this step is to identify one or more SMILES that minimize the number of semantic features that a machine learning model will remember while processing the SMILES.

In the final step, the ClearSMILES are sorted alphanumerically. The first ClearSMILES selected becomes the unique sampled ClearSMILES. This step is implemented to prevent oversampling and improve consistency when working with large batches of randomized Kekulé SMILES.

#### 2.4.3 SELFIES

For the SELFIES we propose two data manipulations to simplify the grammar. The first is to replace overloaded tokens with explicit numerical tokens. Overloaded tokens are the *n* tokens after a branch or ring token, *n* is the digit contained in the ring or branch tokens

##### Algorithm 1

Semantic Memory Map Generation

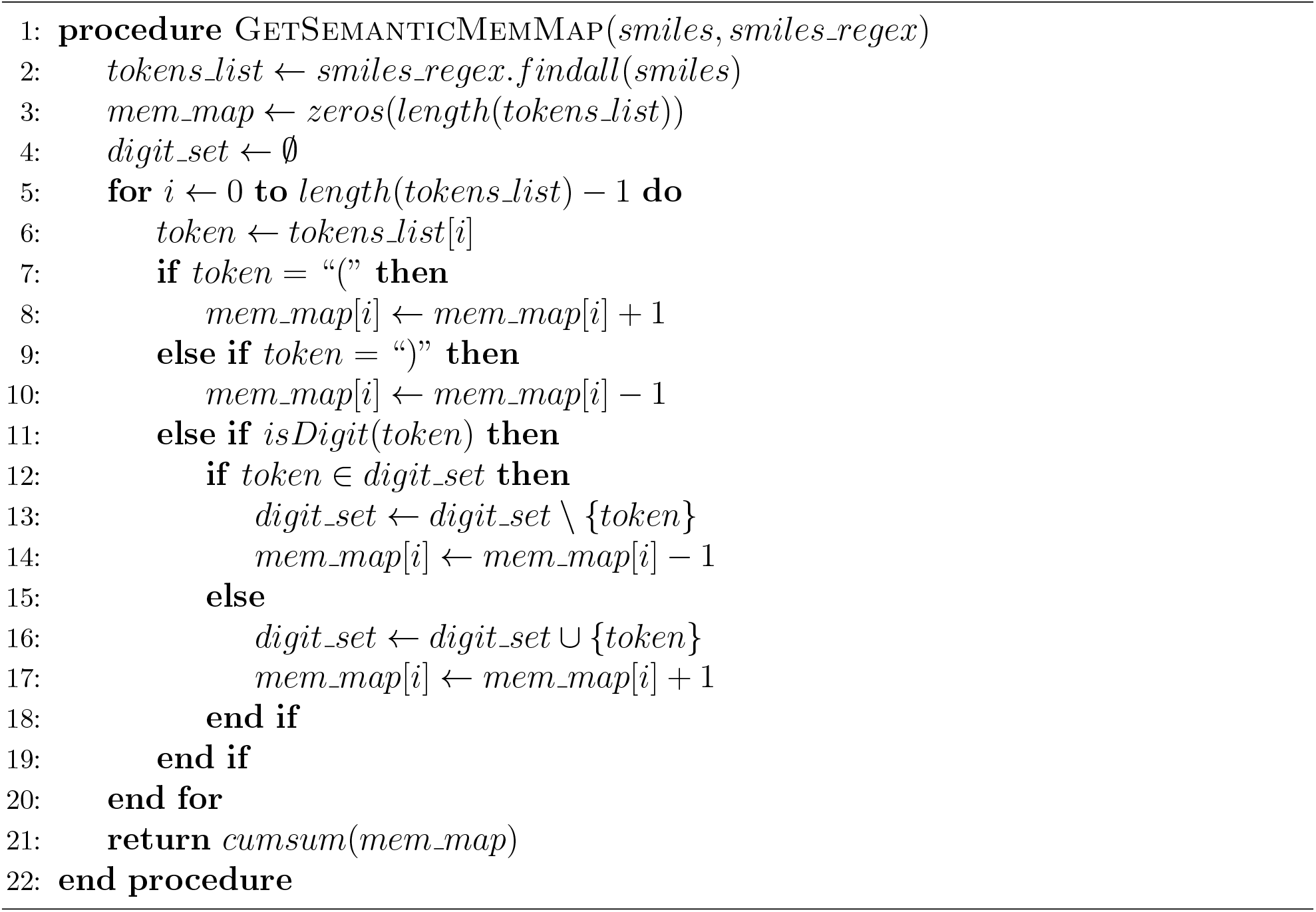

(e.g. [Branch**1**], [Ring**2**]). They defined *N*, the number of tokens in a branch, or the number of atoms that go back to close a ring. To compute *N*, the overloaded tokens are mapped to their index *c*_*k*_ (SI table 1) and used in a hexadecimal system defined by equation 4:

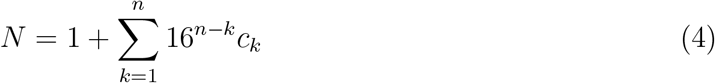

The overloaded tokens are transformed to explicit numerical tokens by replacing their symbols with their corresponding index ([C] to [0]). The second manipulation is to replace the overloaded tokens with a single explicit numerical token indicating the length of the branch or ring, that is, [*N*]. The digit *n* in the branch or ring token preceding the over-loading token (s) is deleted since it no longer has a purpose (Branch2][Ring1][Ring1] to [Branch][17]). The manipulated selfies are always mapped back to their regular counterpart before being decoded to SMILES. This ensures the detection of valence breaks and the syntactical correctness of the SMILES representation.

## 3 Results

### 3.1 regular molecular string representations

#### 3.1.1 Small VAE struggle to emulate molecular string representations

We sampled batches of 300,000 strings from our variational autoencoders (VAEs) trained over 100 epochs. For canonical SMILES-based models, we found validity rates for each batch of samples of roughly 80% (**Table 1**). We found no significant differences in the validity rates when comparing models with 15 and 22 latent dimensions (Supplementary Table **??**).

**Table 1:** viability metrics: validity, novelty,uniqueness for the 300k samples generated by each SMILES based VAE with 22 latent dimensions. All metrics are expressed as percentages.

| Representation | augmentation | Validity | novelty | uniqueness |
| --- | --- | --- | --- | --- |
| SMILES | canonical | 80.75 | **99.57** | 99.92 |
| SMILES | ClearSMILES | **97.80** | 99.13 | 99.92 |

We observed that the string stability of SELFIES generated by non-augmented SELFIES-based VAE is consistently below 50% as shown in Table 2. This implies a partial alteration or loss of information from the original samples during the translation to SMILES, rendering them potentially less suitable as a molecular string representation for a small generative model.

**Table 2:** viability metrics: stability, novelty, uniqueness for the 300k samples generated by each SELFIES based VAE with 22 latent dimensions. All metrics are expressed as percentages.

| Representation | augmentation | Stability | novelty | uniqueness |
| --- | --- | --- | --- | --- |
| SELFIES | regular | 45.42 | 99.91 | 99.96 |
| SELFIES | no overload | 46.42 | **99.93** | **99.98** |
| SELFIES | no hexadecimal | **48.89** | 99.91 | 99.92 |

#### 3.1.2 Identification of Error Origins

Given SMILES and SELFIES are error-prone, it is important to analyze the root causes of errors as a lever to minimize them.

We first looked at the types of errors with the samples produced by the canonical SMILES based VAE. The main source of error is by far aromaticity, but a significant number of errors are also associated with parenthesis, rings, and valence, but not syntax (Figure 2).

**Figure 2:**
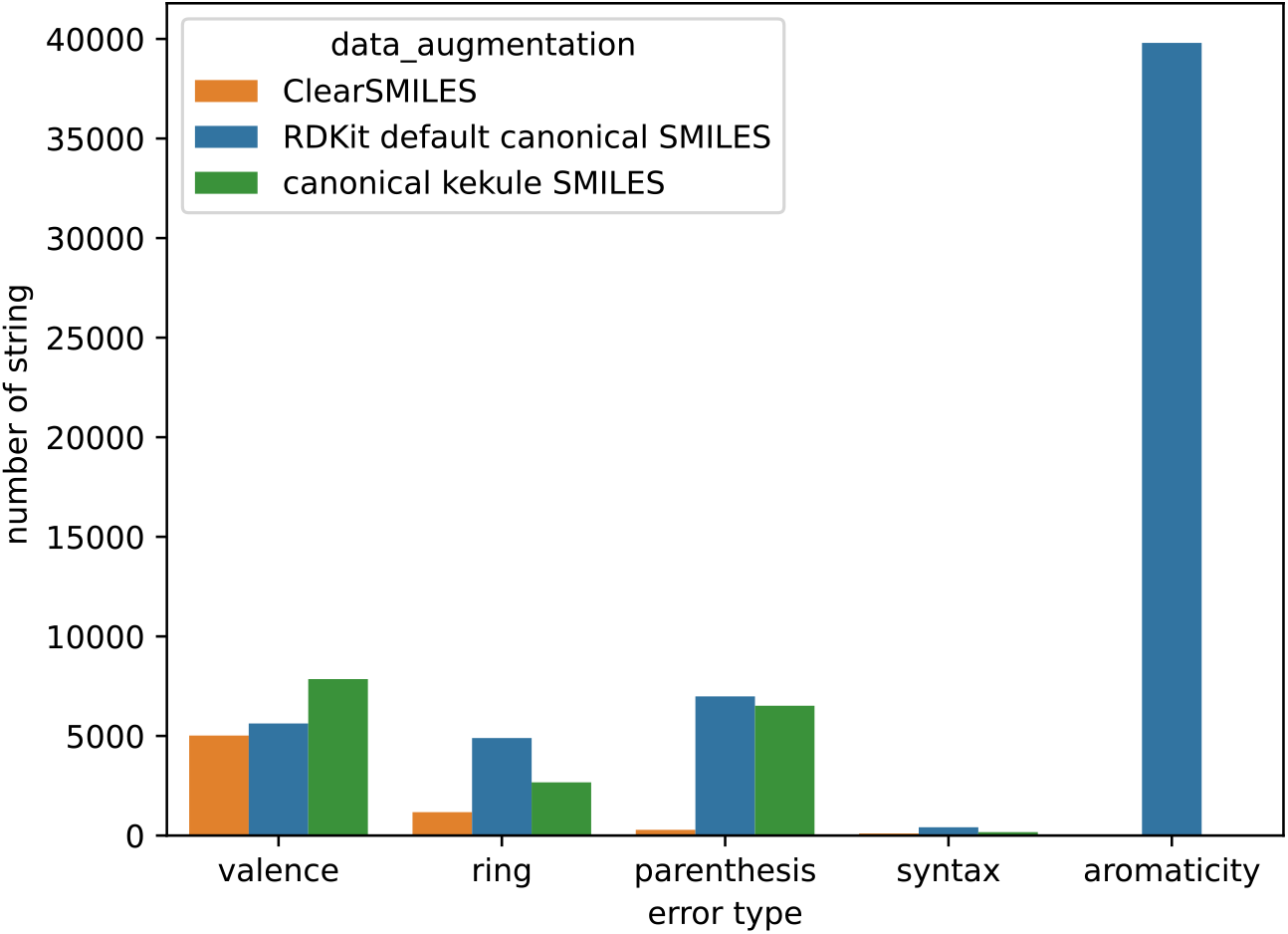
Error of samples generated with canonical SMILES and ClearSMILES based VAEs with 22 latent dimensions

**Figure 3:**
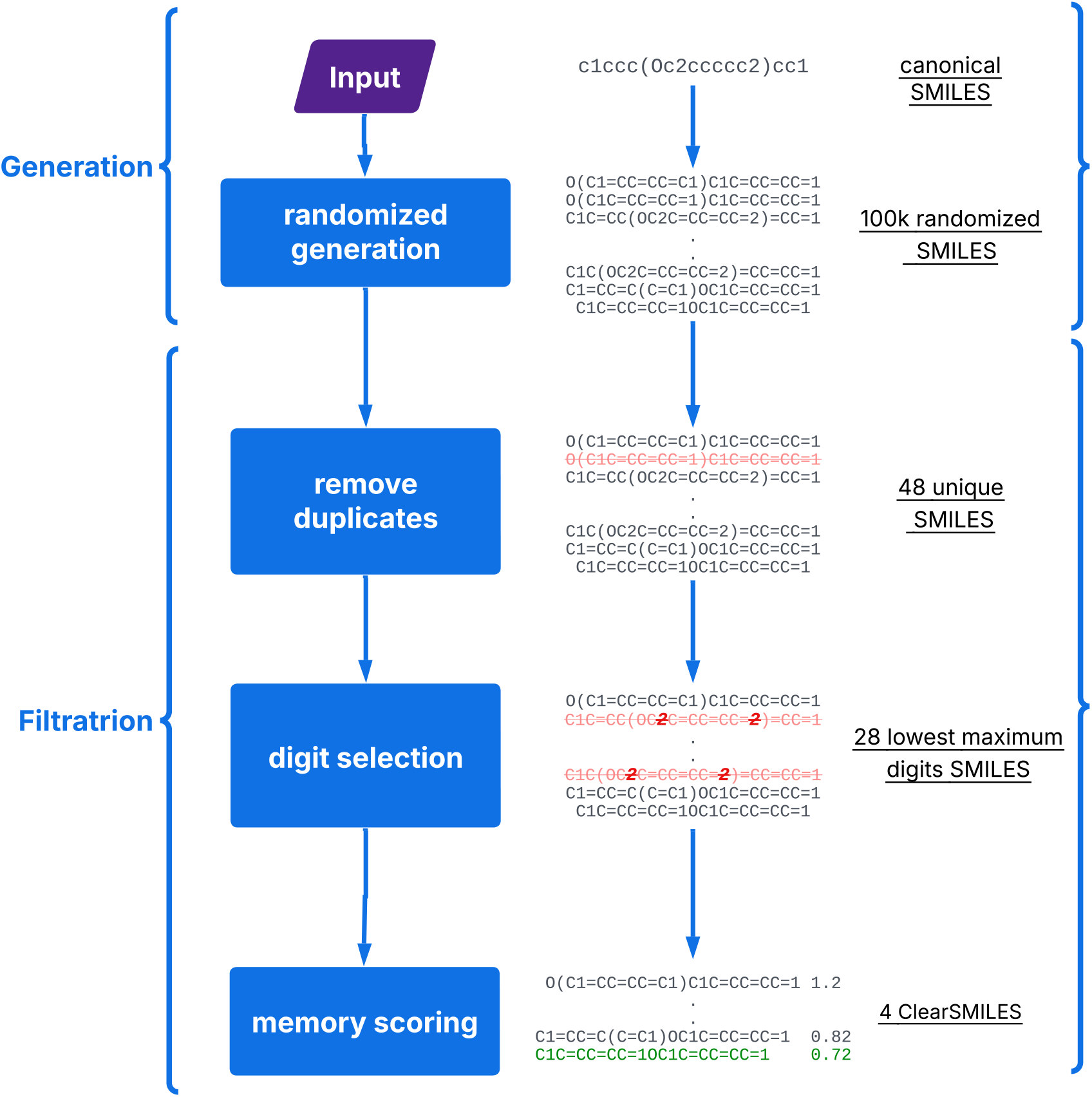
Flowchart of the ClearSMILES pipeline illustrating the generation and filtration of randomized Kekule SMILES, including steps for removing duplicates, selecting minimal-digit SMILES and lowest-memory score SMILES.

We found that there were generally no significant differences in error types between models with 15 and 22 latent dimensions. However, we did notice one notable exception: the occurrence of ring-related errors was 1.84 times more frequent in the SMILES VAE with 15 latent dimensions compared to 22 latent dimensions, as shown in the Supplementary Figure **??**. All the results presented in Figure 2 are consistent with the results described previously in the literature.^26^

Due to the high rate of error caused by poor aromaticity representation and the sensitivity of ring-related error to compression rate in the latent space, we decided to compare the rings present in our samples to those in our training set. We wanted to assess whether the VAE can generate samples with rings that have similar features to those found in the MOSES training set. We selected the number of atoms per ring and the number of rings per molecule as metrics to make our comparison.

We found that only 584 molecules out of 1.6 million present in the MOSES training do not have rings. For molecules with a ring, the number of rings ranged from 1 to 8 and a ring contained 3 to 7 atoms. Most molecules had 2 to 4 rings, mostly 5 or 6 atoms per ring, as illustrated in the Supplementary Figure 10.

We made a contingency table of the previously mentioned ring metrics for both our baseline, the MOSES database, and the samples generated by our VAEs. The MOSES database contains more than one million molecules, whereas batch samples contain only 300,000 molecules. To make a meaningful comparison between our dataset and our samples, we normalize the contingency tables by computing the frequency of each feature. We subtracted the normalized contingency table of MOSES from all normalized contingency tables from the sample batches. The results are presented in Supplementary Figure 11 (a) for samples generated by VAE based on canonical SMILES.

The results for samples generated by the canonical SMILES VAE with 22 latent dimensions show that samples tend to have fewer rings with 6 atoms in molecules with 3 or 4 rings and more rings with 5 atoms in a molecule with 2 rings. Increasing the compression rate of the latent space, from 22 to 15 latent dimensions, slightly widened the already identified disparities between the samples and the characteristics of the baseline ring, as shown in the Supplementary Figure 12. This might indicate a higher rate of failure for molecules with bigger rings and more rings, leading to a skewed distribution of the ring feature where molecules with smaller rings and fewer rings are more prevalent.

We checked the proportion of outlier rings, meaning those with characteristics not found in the MOSES training set, such as rings with more than 7 atoms or molecules containing more than 8 rings. We found that only 0.77% of the rings were outliers in the samples generated by Canonical SMILES VAE with 22 or 15 latent dimensions, indicating that outlier rings are a minor issue.

Considering the above observations, the SMILES should be augmented to limit long-range dependencies between matching tokens and digits, i.e. reducing the token distance between neighboring atoms. This aims to reduce the attention effort required to process an SMILES for the RNNAttn-VAE. The aromaticity of rings should also be explicitly stated in SMILES with an easily recognizable pattern of tokens. To that end, we propose a new data augmentation for SMILES: ClearSMILES (see below).

To ascertain the validity of the SELFIES string stability as a metric, we performed a negative control with the entire MOSES dataset. All SMILES from MOSES were encoded to SELFIES, and a stability assessment was conducted ten times for each SELFIES. Therefore, the stability assessment was repeated 19 million times, and no SELFIES generated from the MOSES dataset failed, indicating that 0% of the MOSES’s SELFIES are unstable.

For unstable SELFIES samples, we computed the normalized token difference between the samples and their regenerated counterpart. We found that in the overwhelming majority of cases unstable SELFIES exhibit some sort of loss of information during their translation to SMILES. We aligned the unstable SELFIES with their regenerated SELFIES using the Needleman-Wunsh algorithm. We found that loss of information is predominantly related to the deletion of branches and rings (see Supplementary Information). We repeated the same analysis of the ring features we performed on the SMILES samples for the SELFIES samples. We found that SELFIES exhibit the same problem as SMILES, leading to a skewed distribution in favor of molecules with smaller and fewer rings. The notable difference is that selfie samples are even further away from the MOSES baseline than SMILES. Also, the percentage of outlier rings is an order of magnitude greater than that of SMILES samples, becoming a non-negligible issue. Our main hypothesis to explain the difference between SMILES and SELFIES is that VAE struggle to emulate the complex hexadecimal encoding using overloaded tokens to define branch and ring length. Thus, SELFIES augmentation should focus on simplifying the encoding of branch and ring length for SELFIES.

### 3.2 Augmented molecular string representation

ClearSMILES has the following objectives: The first aim is to reduce the number of tokens in the vocabulary that are required to represent a full set of SMILES. Thus, a reduction in dimensionality is achieved for the one hot matrix used as input of the VAEs. The second aim is to minimize the number of semantic features open across all tokens in a SMILES. In other words, it means trying to find a solution with the minimum amount of tokens between matching digits and matching tokens, while also minimizing the number of rings or SMILES concurrently open. In a perfect theoretical case, the number of tokens between matching parentheses or digits is very small, and all branches and rings are closed before another one begins. This will limit the attention effort a deep learning model or a machine learning model has to make to process a SMILES. The different steps of the ClearSMILES pipeline are detailed in the Methods section and in Figure 3

For SELFIES we introduced two data augmentation, the first is to remove the overloading for the hexadecimal encoding of branch and ring length, the second data augmentation is to remove the hexadecimal entirely, and replace by a single non-overloaded token. For more details, please refer to the Methods section.

#### 3.2.1 Analysis of augmented string properties

We first tested the relationship between the number of randomized SMILES and the number of unique SMILES using 1000 iterations of 1000 randomized SMILES for approximately 1% of MOSES training set. We counted the cumulative set of unique SMIELS yielded by each iteration. If the size of the cumulative set did not grow for 10 iterations, the generation was prematurely stopped. We found that the overwhelming majority (84.57%) of molecules had reached a plateau before or at 100 iterations, meaning after a cumulative number of 100,000 randomized SMILES generated. Thus, we decided to choose 100,000 randomized SMILES as the number of random SMILES to be generated in the ClearSMILES pipeline, as it can also be computed in a reasonable time.

We applied the ClearSMILES data augmentation procedure to the 1.9 million SMILES originating from the combined MOSES training, test, and scaffold sets. For approximately 96% of molecules it takes less than 12 seconds, with a majority between 5 to 10 seconds. For approximately 4% of the data, it takes between 15 and 40 seconds to create a ClearSMILES from a SMILES. This process yielded a total of 6.9 million ClearSMILES, each having a varying number of equivalent solutions, ranging from 1 to 128. Half of the ClearSMILES generated had two or less solutions, with 80% of them producing four or fewer equivalent solutions, as shown in Supplementary Table 4. The highest decile encompassed the widest range, ranging from 8 to 128 equivalent solutions.

To prevent oversampling of certain molecules and ensure some level of consistency, ClearSMILES are sorted alphanumerically using the default python sorting algorithm, and only the first ClearSMILES is kept. The retained ClearSMILES will later be referred to as the sampled ClearSMILES.

Our initial comparison focused on the number of tokens per string for canonical SMILES and sampled ClearSMILES, showing a change in the distribution of token lengths between the two, as illustrated in the Supplementary Figure 8. Specifically, sampled ClearSMILES tend to have more tokens per string compared to their canonical counterparts, as summarized in the Supplementary Table 6.

However, it is worth noting that the difference is relatively smaller when considering the longest ClearSMILES and canonical SMILES, i.e. the SMILES with the highest number of tokens for both categories. We found that the longest ClearSMILES is only 9.25% longer than the longest canonical SMILES. This is important, as the size of longest SMILES is equal to the number of row in the one-hot encoded matrix used as input to the VAE. This means that the increase in the number of rows for ClearSMILES’s one hot encoded matrices is fairly limited compared to the canonical SMILES’es one-hot encoded matrices. We then performed a comparative analysis between the sampled ClearSMILES and the canonical SMILES, with a specific focus on their memory scores, branches, and rings.

The memory scores obtained from sampled ClearSMILES and the canonical SMILES generated by RDKIT exhibit significantly different distributions, as depicted in **Figure 4 (b)**. The memory scores for sampled ClearSMILES form a Gaussian-like distribution centered around 0.86 with a small standard-deviation of 0.13. In contrast, canonical SMILES memory scores have a distribution that strays further from a Gaussian distribution with a standard deviation of 0.66. Also, the average memory score for ClearSMILES is 0.86, approximately half of the average memory score of 1.72 for canonical SMILES.

**Figure 4:**
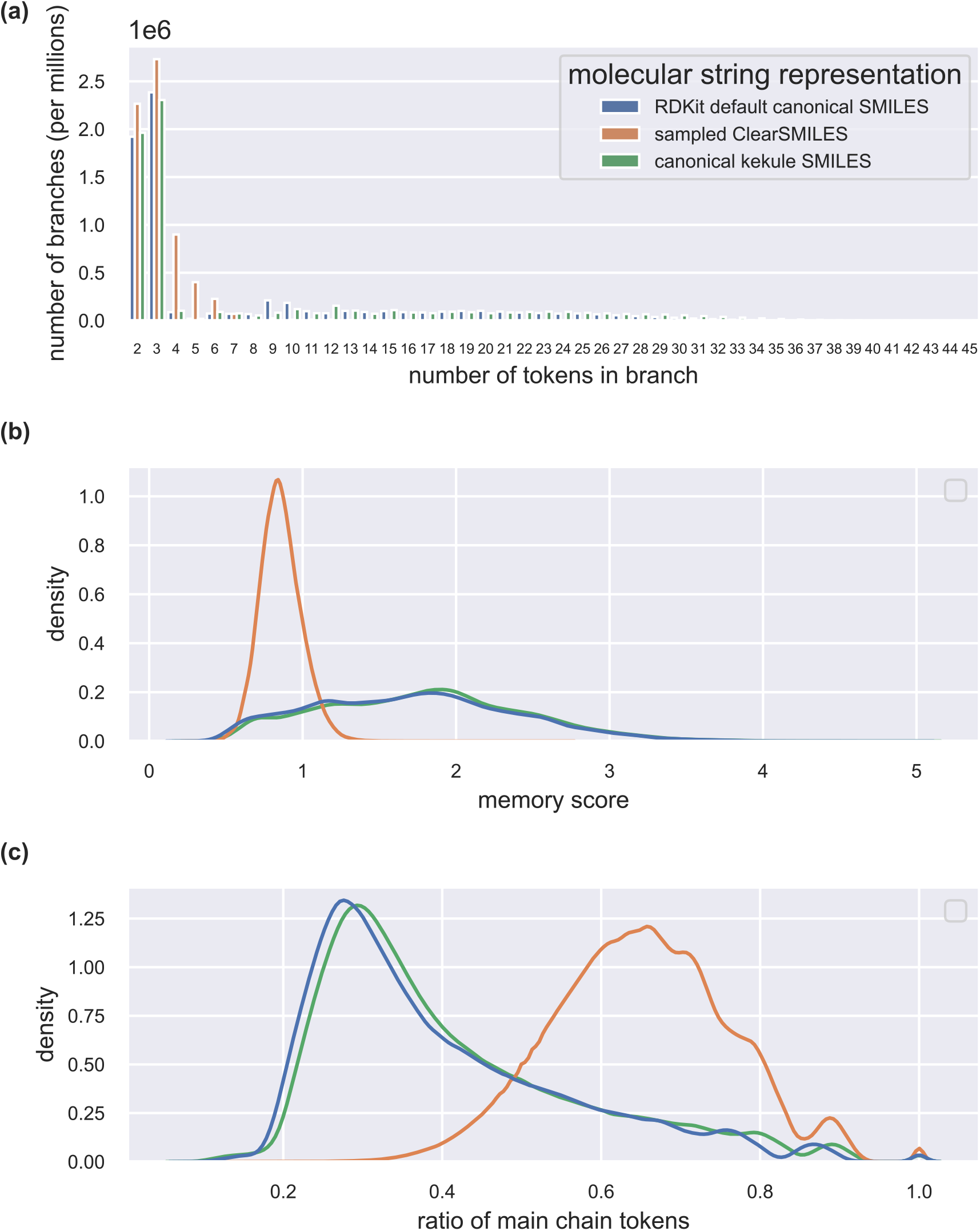
Histogram of the branch token distribution (a), kernel density estimation (kde) of the memory score (b) and the ratio of main chains tokens for the MOSES database: canonical SMILES (blue) and ClearSMILES (orange)

Quantile analysis of memory scores reveals that around 90% of sampled ClearSMILES have a memory score below or equal to 1, while fewer than 20% of canonical SMILES exhibit a memory score below or equal to 1. This indicates that for an overwhelming majority of sampled ClearSMILES can be processed with a low attention effort by a neural network, whereas only a few canonical SMILES can.

We computed for each molecule the difference between the number of branches in the sampled ClearSMILES and the number of branches in the corresponding canonical SMILES, which we refer to as Δ*branches* (SI Table 5). In roughly 75% of the cases, there was no variation in the number of branches between a ClearSMILES sample and its corresponding canonical SMILES. There are slightly more sampled ClearSMILES with more branches than sampled ClearSMILES with less branches compared to their canonical SMILES counterpart. Overall, our findings indicate that sampled ClearSMILES exhibit very little alteration in the distribution of the number of branches compared to that of canonical SMILES.

However, a substantial difference emerges in the distribution of branch sizes between canonical SMILES and sampled ClearSMILES, as depicted in **Figure 4 (a)**. Sampled ClearSMILES tend to exhibit a greater prevalence of branches with sizes falling within the range of 2 to 6 tokens. Notably, there is a remarkable 52-fold reduction in the occurrence of branches strictly longer than 10 tokens in sampled ClearSMILES compared to canonical SMILES.

After analyzing the distribution of branches in Sampled ClearSMILES and canonical SMILES, we looked at the differences in terms of ring representation in strings between ClearSMILES and canonical SMILES. Because the number of rings is an inherent property of a given molecule, it remains invariant between canonical SMILES and sampled ClearSMILES and therefore was not studied.

Note that the distances between pair digits that represent a ring closure can vary depending on the path taken to traverse the molecule to generate a SMILES. If a ring closure is open while another is still open, the digit representing the new ring closure is obtained by increasing the digit of the ring still open by one. When a ring is closed, the digit used to represent it can be reused for a new ring opening. Therefore, the maximum number of digits required to describe all rings can also fluctuate on the basis of the separation degree between the rings.

We computed the contingency table for the number of branches using the following two criteria: digit used in each pair representing a ring closure and the number of tokens between the paired digits. Due to the substantial difference in scale between each category, we used a logarithmic base 10 scale to enhance readability, as illustrated in Figure 5.

**Figure 5:**
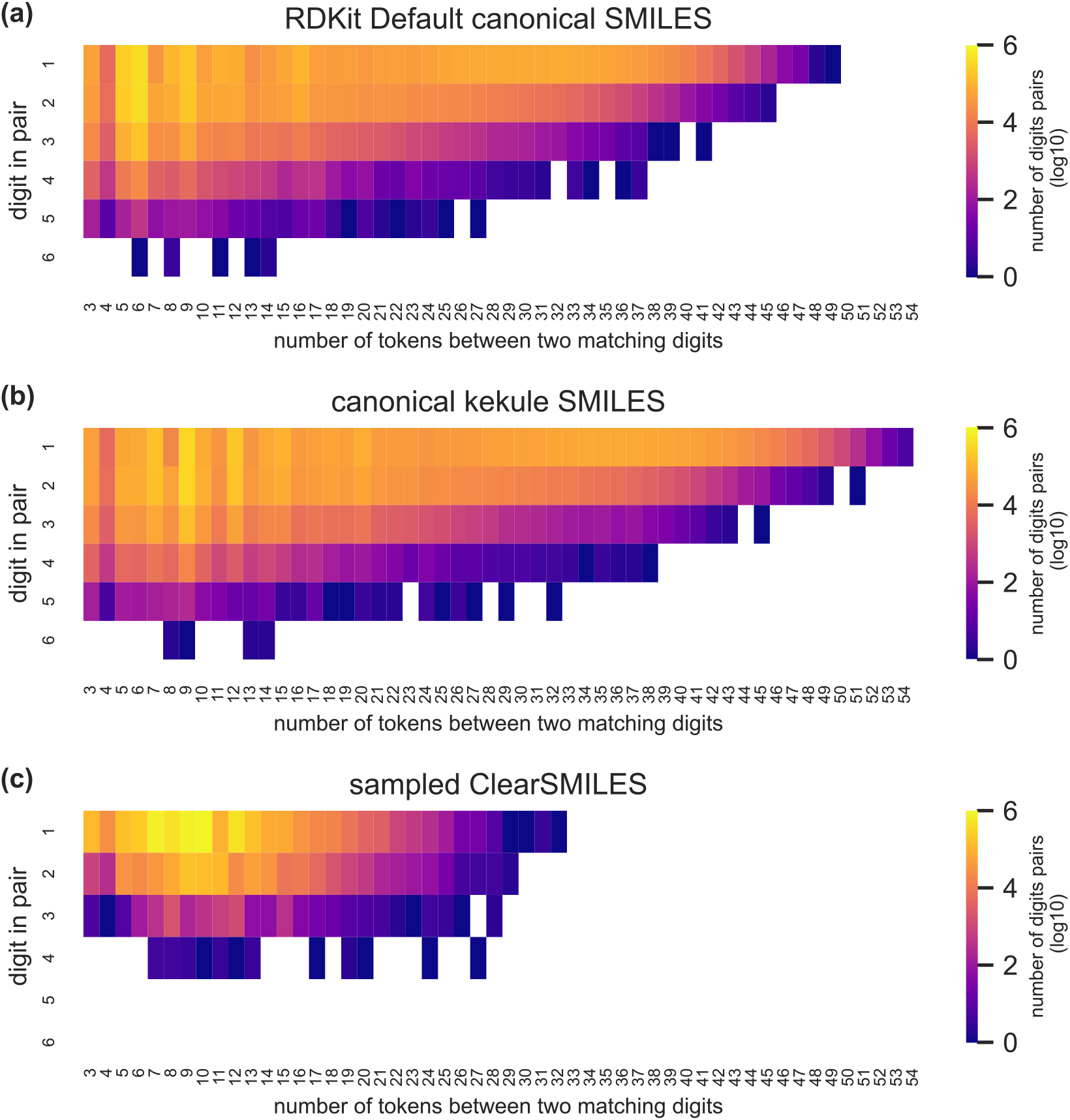
Heatmap illustrating the contingency table of paired digits in SMILES-based molecular string representation. A base-10 logarithmic scale is applied to enhance visibility, given the large differences in scale between categories. White spaces represent categories with zero values

Sampled ClearSMILES require significantly fewer digits than canonical SMILES to represent the same molecules, as depicted in Figure 5. In fact, just two digits are sufficient to describe 99.72% of the sampled ClearSMILES, a stark contrast to canonical SMILES, where only 74.27% can be represented with two digits.

Moreover, sampled ClearSMILES exhibit a notable reduction in the occurrence of paired digits with substantial gaps between them compared to canonical SMILES, as illustrated in Figure 5. There is a remarkable 4 order of magnitude decrease in the number of paired digits with gaps exceeding 25 tokens between them with sampled ClearSMILES compared to canonical SMILES. However, paired digits with gaps of less than 16 tokens are more frequent in canonical SMILES. For smaller gap values between pair digits, Sampled ClearSMILES tends to have slightly more gap because the kekule representation in ClearSMILES uses a mixture of uppercase atom symbol (aliphatic) coupled with double bonds to represent aromaticity. In canonical SMILES aromaticity is simply represented by lowercase atom symbols. Thus, representing aromatic rings in ClearSMILES will always use more tokens than in canonical SMILES.

We then studied the graph complexity associated with our molecular string representation. Evaluating the complexity of a string representation presents a non-trivial challenge. The most effective proxy measure we identified was quantifying the number of tokens within the primary chain of the string. A higher token count in the main chain indicates a greater linearity in the string graph. The increased linearity in the graph, in turn, corresponds to greater simplicity in the graph structure.

Sampled ClearSMILES and canonical SMILES exhibit distinct distributions in terms of token length. To make a meaningful comparison, we calculated the ratio between the number of tokens in the main chain and the total number of tokens. This approach reveals that sampled ClearSMILES display a more uniform distribution, generally yielding higher ratios compared to canonical SMILES, as depicted in **Figure 4 (c)**.

For training purposes, we kept only the sampled ClearSMILES where the highest digit value used was 2 or lower, indicating up to two rings. This led to the exclusion of approximately 0.27% of the initial dataset, which predominantly contained molecules with multiple rings that share two bonds, such as the Adamantyl group, found in half of the excluded molecules. Additionally, we removed the longest strings based on token count, setting a maximum token limit of 58. The original MOSES data set train and test split was otherwise maintained.

#### 3.2.2 Influence on Vocabulary and One-Hot Encoding

In natural language processing, vocabulary refers to the collection of tokens or words required to describe the entirety of the corpus, which, in this case, is the MOSES dataset. It is important to note that vocabulary is representation-specific, meaning that it will differ between SMILES, SELFIES, and their augmented counterparts. ClearSMILES have a vocabulary length of 16 tokens, with the following tokens excluded: Aromatic tokens: ‘[nH]’,’c’,’n’, ‘o’, ‘s’; Bond token: ‘-’ and Digit token: ‘3’,’4’,’5’,’6’. This results in a 30% reduction in vocabulary size with ClearSMILES compared to canonical SMILES vocabulary (cf. SI).

The removal of overloading from the SELFIES grammar resulted in the addition of 16 tokens and the removal of 1 token, increasing the total from 25 tokens to 40 tokens. This represents a 60% increase in the number of tokens. The newly added tokens are all numerical (0-15).

The removal of the hexadecimal system in SELFIES led to the inclusion of 45 new tokens and the removal of 11 tokens, resulting in a total token count increase from 25 to 59. This means a 2.36-fold increase in the number of tokens.

In the vocabulary of all molecular string representations, we’ve added a few special tokens: ’<start>’ to mark the beginning of a string, ’<end >’ to signal its end, and ’_’ to pad strings to a uniform maximum length.

We conducted a comparison of the dimensions of the one-hot matrices. These matrices are characterized by dimensions of *n × m, n* represents the maximum number of tokens per string, and *m* corresponds to the vocabulary size. A comprehensive summary of the one-hot matrix dimensions can be found in **Table 3**.

**Table 3:** Summary of one-hot encoded matrix dimensions for each molecular string representation. The resizing factor represents the ratio of the number of dimensions in the current string representation to the number of dimensions in the original string representation.

| String Molecular Representation | n | m | Resizing Factor |
| --- | --- | --- | --- |
| SMILES | 56 | 26 | 1 |
| ClearSMILES | 58 | 19 | 0.78 |
| SELFIES | 57 | 28 | 1 |
| SELFIES no overload | 57 | 43 | 1.53 |
| SELFIES no hexadecimal | 52 | 62 | 2.02 |

Our analysis revealed that ClearSMILES achieved a 22% reduction in the size of the one-hot matrices when compared to SMILES. Augmented SELFIES exhibited larger one-hot matrices, showing a 53% increase for SELFIES without overload and a 2.02-fold increase for SELFIES without the hexadecimal system. For simplicity sake, the n of all matrices were set to the highest value of n, which is 58.

#### 3.2.3 Analysis of ClearSMILES performance

We sampled batches of 300,000 strings from our ClearSMILES-based VAEs which showed a notable change compared to those based on canonical SMILES. This shift led to a substantial increase of around 18 points in validity rates, i.e. the number of invalid molecules is decreased by roughly an order of magnitude (Table 1). There is a slight decrease of one percent point in the validity rates when going from 22 to 15 latent dimensions for samples generated by the ClearSMILES VAE as shown in the Supplementary Table **??**.

The samples generated by ClearSMILES-based VAEs demonstrate a substantial reduction in the number of errors in all categories, except for valence errors. In this specific category, samples generated by ClearSMILES VAE with 15 latent dimensions exhibit a slight increase in the number of errors. Samples generated by the ClearSMILES VAE with 22 latent dimensions show a slightly lower number of errors compared to samples generated by the canonical SMILES VAE. (Figure 2).

It is important to note that ClearSMILES lacks the lowercase aromaticity representation, which means that RDKit cannot classify any error as aromaticity errors in ClearSMILES samples. Any erroneous mixture of aliphatic atoms with double bonds, which explicitly represents the conjugated system of aromatic rings in ClearSMILES, is categorized as a valence error instead of an aromaticity error. Even taking this fact into account, it is evident that ClearSMILES enables VAE to drastically reduce the number of valence and aromatic errors as their sum is much smaller than the sum of those errors with samples generated from the canonical SMILES VAE. A possible explanation to why there is a higher amount of valence error for samples generated with a ClearSMILES VAE with 15 latent dimensions can be that there is an increase in the aromaticity error that is mislabelled as a valence error by RDKit.

To further our analysis, we proceeded to compute the contingency table of ring features for the samples generated by the ClearSMILES VAE with 22 and 15 latent dimensions are shown in the Supplementary Figure 11 (b) and the Supplementary Figure 12 (b). We found that samples generated by a ClearSMILES VAE with 22 latent dimensions have ring feature closer to the MOSES baseline than the samples generated by a canonical SMILES VAE with also 22 latent dimensions. However, this comparison does not hold for samples generated by VAE with 15 latent dimensions, where canonical SMILES outperforms ClearSMILES on this metric. Canonical SMILES samples generated by a VAE with 15 or 22 latent dimensions do not exhibit a significant difference in ring feature. However, for ClearSMILES samples there is a moderate yet noticeable stray from the MOSES ring feature distribution when the compression rate increases from 22 to 15 latent dimensions. This difference could be explained by the fact that the lowercase representation of aromaticity in canonical SMILES is more compact than the kekulé representation in ClearSMILES which could be more resilient to compression in the latent space.

For rings with outlier length, i.e. rings with more than 7 atoms, ClearSMILES-based model does not generate rings with more than 13 atoms per ring. The canonical SMILES-based model generated samples with up to 17 atoms per ring. Thus, the use of ClearSMILES limits the degree of ring ‘aberrancy’ without completely eliminating it. The percentages of outlier rings for samples generated by 15 and 22 latent dimensions ClearSMILES VAE are 0.38% and 0.29%, which is approximately half the rate of outlier rings for samples generated by either canonical SMILES VAE(s).

#### 3.2.4 Analysis of augmented SELFIES performance

The data augmentation proposed for SELFIES did not result in a substantial enhancement of string stability rates, with the most significant improvement being a maximum of a 5-point increase observed between regular SELFIES and no-overload SELFIES when using a VAE with 15 latent dimensions. Interestingly, the transition from VAEs with 22 latent dimensions to those with 15 latent dimensions showed a marginal but consistent improvement in string stability rates across all data augmentation procedures and regular SELFIES.

We analyzed the token loss of each batch of 300K samples generated by the augmented SELFIES VAE. We computed the details of the distribution using the kernel density estimation (see Supplementary Information). We found that there is no distinct change in the distribution of the token loss for augmented SELFIES. We categorize the token loss as loss, gain or unchanged, as illustrated in the Supplementary Table 2. Around 91% of the unstable augmented SELFIES have a net loss of tokens. The rest of the unstable augmented SELFIES experience either a gain in token or no change in the token length.

The rings feature of samples were analyzed with contingency as before, showing that for samples generated by augmented SELFIES VAE with 22 latent dimensions, the ring features stray further from the MOSES baseline than the samples generated by the SELFIES VAE.

However, it is the other way around with samples generated by VAE with 15 dimensions, although the improvement is marginal. In general, the percentages of aberrant rings for augmented samples remain in the same order of magnitude as the percentages of the original SELFIES samples with a range of 6% to 8%.

### 3.3 Fidelity Assessment of all models

We conducted a fidelity assessment of compounds using various metrics, including quantitative estimate of drug-likeness (QED), synthetic accessibility estimation (SA), molecular weight(MW) and topological polar surface area (TPSA). Kernel density estimation (KDE) was computed for each metric, as depicted in Figure 6. Furthermore, we calculated the Wasserstein distance for each metric between all models and the MOSES train set baseline to ensure a fair comparison. The results for models with 22 latent dimensions are presented in Table 4. The results for all models are available in the Supplementary Table 8.

**Table 4:** Wasserstein distance between the quantitative metrics (Molecular Weights, QED, SA, TPSA) and ring-related descriptors of the 300k random samples from the MOSES training set and each batch of 300K samples generated by VAE(s) with 22 latent dimension.

| Model | TPSA | MolWeight | QED | SA |
| --- | --- | --- | --- | --- |
| SELFIES no hexadecimal | 7.798 | **6.805** | 0.115 | 1.316 |
| SELFIES no overload | 10.094 | 6.693 | 0.141 | 1.315 |
| regular SELFIES | 4.363 | 7.017 | 0.112 | 1.258 |
| canonical SMILES | 4.716 | 7.149 | 0.041 | 0.430 |
| ClearSMILES | **4.304** | 7.344 | **0.022** | **0.345** |

**Figure 6:**
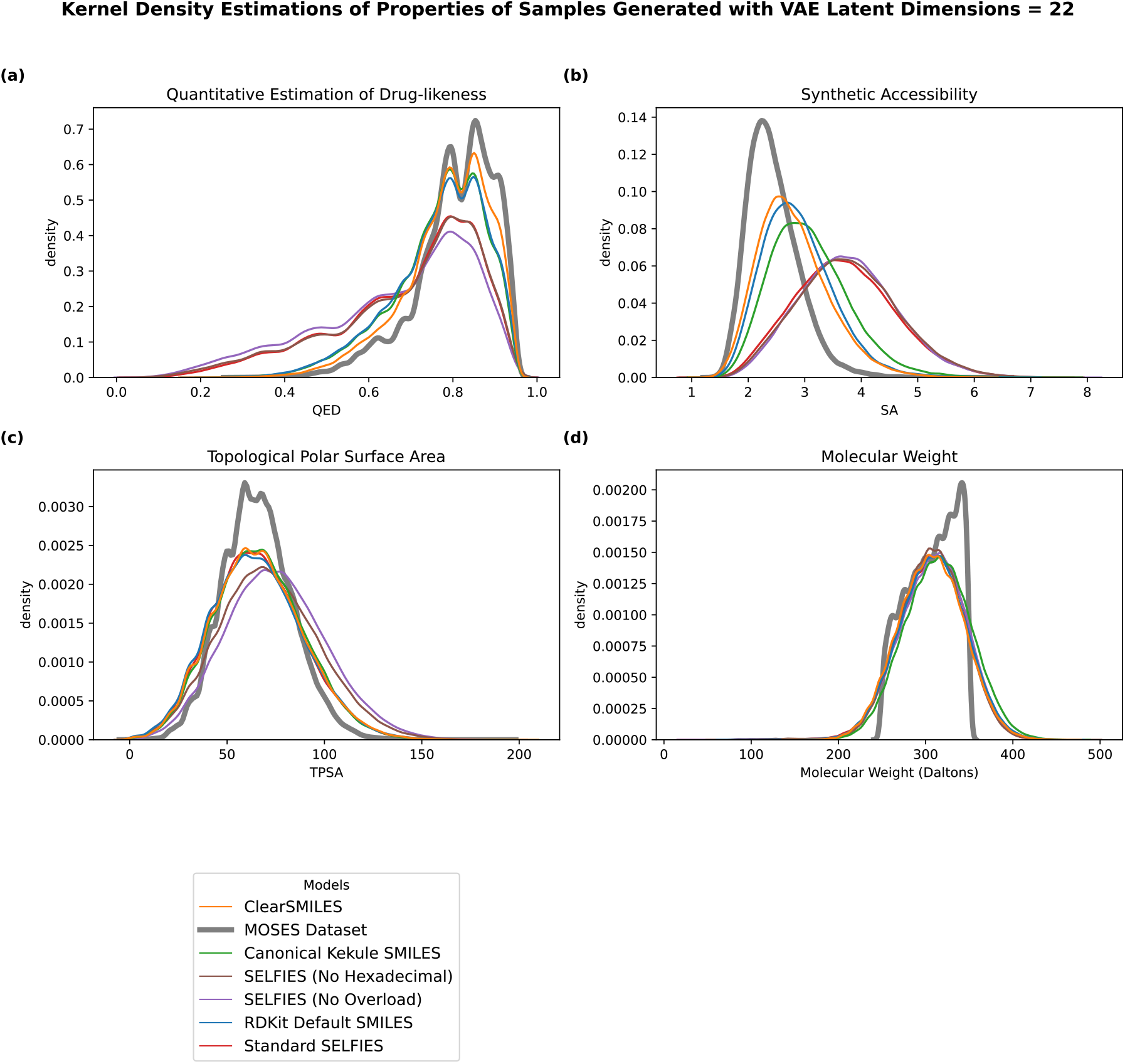
Assessment of various metrics for all valid samples generated by all models: (a) QED, (b) SA, (c) TPSA, (d) molecular weight

Our analysis revealed that for QED, as illustrated in Figure 6 (a), SMILES-based models generate samples that consistently fit better to the training set compared to the SELFIES-based models generated. SELFIES-generated samples tend to exhibit lower QED values, indicating molecules that are less drug-like. This discrepancy is particularly evident for QED values below 0.6, with models based on SELFIES producing a higher number of samples in this category than models based on SMILES. Table 4 highlights that the Wasserstein distance for QED between MOSES and samples is approximately three times lower for SMILES samples compared to SELFIES samples, confirming that SMILES samples align more closely with the MOSES baseline. Notably, the samples generated by the 22 latent dimension VAE using ClearSMILES as its string molecular representation achieve the lowest Wasserstein distance with a very satisfying fit.

The density kernel estimation for the synthetic accessibility estimation score (SA), shown in Figure 6 (b), indicates a clear difference between the distribution of samples and the MOSES baseline. SELFIES samples consistently underperform in terms of synthetic accessibility, with SMILES-based models exhibiting a distribution closer to the MOSES baseline than SELFIES-based models. The difference in Wasserstein distance further confirms this trend, with SMILES samples showing approximately 2 to 5 times lower values than SELFIES samples.

The TPSA distributions of the samples, in all representations, generally align well with the MOSES baseline, as shown in Figure 6 (c). Wasserstein metrics suggest comparable performance between SMILES-based and regular SELFIES samples, with a slightly lower distance for regular SELFIES. However, augmented SELFIES samples deviate further from the baseline in terms of TPSA, showing a 2-fold increase in the Wasserstein distance compared to regular SELFIES.

The molecular weight distributions of the samples, considering all representations, also align well with the MOSES baseline, as illustrated in Figure 6 (d). Wasserstein distance analysis indicates that the augmented SELFIES slightly outperforms other representations in terms of proximity to the baseline. Interestingly, there seems to be a slight positive effect on the Wasserstein distances by increasing the compression rate of the VAE, transitioning from 22 latent dimensions to 15 dimensions, except for samples generated by regular SELFIES VAEs as shown in the Supplementary Table 8.

## 4 Discussion

In this study, our results suggest that for small generative models, canonical SMILES and SELFIES may not be sufficiently robust string molecular representations to be used as is. The error analysis of samples generated by a Canonical SMILES based VAE revealed multiple difficulties encountered by a small generative model. The first difficulty is in accurately capturing abstract chemical properties, such as aromaticity and ring features. The lowercase representation of aromaticity in RDKit canonical SMILES is probably too abstract for a simple RNN to relate to its aliphatic counterparts. We speculate that the one-hot encoded matrix lacks an apparent relationship between aromatic and aliphatic groups due to their encoding in separate columns, unlike in string representations where a relationship is apparent between uppercase and lowercase symbols. The description of the connectivity and electronic system is also abstracted away, which means that VAEs using canonical SMILES must infer a complex set of rules about connectivity between aromatic atoms from little to no information. In Kekule SMILES, a specific mesomer is picked, which allows the description of an aromatic system as a combination of aliphatic tokens and double bonds tokens. This removes the need to establish a relationship between aromatic symbols and aliphatic symbols as aromaticity can be described without the use of lowercase symbols.The combination of tokens introduced by the Kekule representation likely forms a repetitive pattern that the VAE can recognize, aiding in the differentiation from aliphatic rings.

The results suggest that Recurrent Neural Networks (RNNs) in the VAE are able to grasp the general syntax of SMILES but struggle to consistently keep track of parentheses (branches), digits (rings), and valence across a long SMILES. This is related to a known problem with RNN called the vanishing gradient problem, wherein training on a long sequence of tokens weakens the gradient used for back-propagation, making it increasingly difficult to properly capture long-term dependencies. ^27,28^ One of the most common ways to deal with this problem is self-attention,^29,30^ a mechanism that aims to mimic human attention. However, considering the previously mentioned results, the single head of attention in the RNNAttn-VAE does not seem to be enough to fully compensate for the vanishing gradient problem. Furthermore, it seems that RNN in our VAE struggle even more with abstract chemical properties such as aromaticity. The Canonization algorithm for SMILES aims to provide a consistent way to find a single-atomic ranking to generate a unique SMILES per molecule. It does not seek to find a solution that minimizes or limits the number of long-term dependencies. To mitigate previously mentioned issues, we introduced ClearSMILES a stochastic data augmentation procedure that takes advantage of the uniquess problem. Many SMILES can represent a same molecule because each unique combination of starting and path can generate a different SMILES. However, not all combinations of starting/path for SMILES are equivalent. The length and the number of long range dependencies in a SMILES are intrinsically tied to the combination used to make the SMILES. You can also choose the level of the SMILES representation, you can include geometric information such as stereoisomerism, explicitly represent hydrogen atoms, and single bonds. Therefore, it can be said that: ‘SMILES are equal, but some are more equal than others’. The core principle of ClearSMILES is to choose the SMILES for which the path and starting point minimize long-term dependencies and choose a level of representation that simplifies learning the distribution of complex concept such as aromaticity. ClearSMILES explicitly takes into account the long-term dependencies in a SMILES with the memory score, a heuristic serving as a proxy for the number and arrangement of open semantic features (ring closure or branch) across a SMILES. The lower the memory score, the fewer concurrently open semantic features. ClearSMILES identifies unique randomized Kekulé SMILES with minimal digits for ring closures and the lowest memory score. Fewer digits decrease the One-Hot Encoded Matrix dimensionality for VAE input, further reduced by replacing the lowercase aromatic tokens by a mixture of already used tokens. SMILES with low memory scores reduce long-range dependencies by minimizing tokens between paired parentheses and digits, leading to lower graph complexity. Removing aromatic tokens and minimizing digits for SMILES reduces vocabulary by 30%, resulting in a one-hot encoded matrix that is 22% smaller than that for canonical SMILES.

ClearSMILES reduces the generation of invalid SMILES from our VAE by an order of magnitude, increasing the validity rate from 80% to almost 98%. Samples generated using ClearSMILES had a druglikeness (QED), synthetic accessibility (SA) and Topological Polar Surface Area (TPSA) closer to the MOSES training baseline than the one generated using canonical SMILES. Therefore, although samples generated by Canonical SMILES slightly outperform ClearSMILES in terms of molecular weight, ClearSMILES largely outperforms SMILES in terms of both validity and fidelity.

We found that most samples generated using SELFIES as a molecular string representation experienced instability. This means that they could not be translated into SMILES without some loss or alteration of information. They also struggled more than SMILES-based representations in faithfully reproducing ring features. SELFIES produced about 7% outlier rings, compared to less than 0.8% for SMILES. This is likely due to the significantly more complex syntax of SELFIES, which makes it more difficult for a small generative model to properly emulate the ring features.

VAEs using SELFIES struggle to faithfully reproduce critical dataset properties such as druglikeness and synthetic accessibility. The emphasis of the SELFIES algorithm on achieving 100% validity results in information loss, particularly in the deletion of ring closures and branches, leading to “malformed” molecules with excessively large rings or improbably long chains. The comparison of Wassertein distances between stable and unstable SELFIES indicates that unstable SELFIES contribute significantly to the poor performance of sampled SELFIES. This is consistent with the fact that unstable SELFIES lose part of their information when translated to SMILES due to an erroneous token arrangement. This is consistent with the findings of the Texas SELFIES,^31^ where by authorizing the decoding of SMILES with valences rule break, the author was able to remove low quality samples.The data augmentation proposed for SELFIES in this study provided only a limited increase in string stability and did not address issues such as outlier rings or the chemical quality of the samples. The primary limitation appears to be the curse of dimensionality, linked to the substantial increase in vocabulary size associated with replacing overloaded tokens. Future work on SELFIES could explore ways to limit the number of tokens used to describe branch lengths and positions without resorting to overloading.

The number of latent dimensions in the original paper of RNNAttn-VAE^7^ is 128 or 256 latent. The author’s choice is interesting because it diverges significantly from the VAE principle of compressing data in the latent space to retain only essential input features. Taking into account the length (*n*) of the one-hot input matrices of the PubChem subset (*n* = 128) or their ZINC subset (*n* ≈ 54) we can see that there is little to no compression but rather a decompression. Even considering the size of the vocabulary which is somewhat small, roughly 30 unique tokens, mapping sparse binary data to a continuous representation following a standard normal distribution appears trivial for a RNNAttn-VAE. Therefore, we assumed that the RNNAttn-VAE could in fact learn relevant features of data at a much higher rate of compression. In that spirit, we chose to start with 22 latent dimensions to have a little bit more than a twofold compression compared to our input size (*n* = 56 − 59).

To confirm this hypothesis, we reused the Shannon Information Entropy measurement for each latent dimension of the latent space introduced by Dollar et al.^7^ Shannon entropy allows us to detect the repartition of information in the latent space. We show that the data repartition is well distributed for all SMILES-based models, as shown in SI Figure 15.

## 5 Conclusion

In summary, our study successfully improved the reliability of SMILES by implementing a new data augmentation technique called ClearSMILES. This approach increased the validity rate of the samples by approximately 18 percentage points and improved the ability of a small variational autoencoders (VAE) to generate samples that faithfully capture the chemical properties of compounds present in training sets. ClearSMILES achieves a compact data representation, limiting the size of the one-hot encoding matrix compared to that of the canonical SMILES for identical molecules.

Although the SELFIES representation ensures that 100% of SELFIES can be translated to SMILES with a correct valence, it leads to SELFIES instability, where information loss or alteration occurs when a sequence of SELFIES tokens with incorrect valence is altered or removed when translated to SMILES. Unfortunately, more than half of the generated SELFIES-based samples exhibit this instability, primarily characterized by the deletion of ring information during translation, resulting in a notable outlier ring rate, with rings containing up to 22 atoms. Furthermore, SELFIES samples less faithfully reproduce critical chemical properties found in the training set compared to SMILES. This is particularly true for unstable SELFIES, in terms of their estimated drug-likeness and synthetic accessibility. As a result, we conclude that SELFIES, in their current form, may not be the most suitable representation for small generative models. They likely require more complex transformer-based models to better understand their complex syntax and reduce instability issues.

## 6 Authors Contribution

Zoë Wefers: Conceptualization (simplified SELFIES approach)Software (RNN-AttnVAE code adaptation), Writing - Original Draft and Investigation Etienne Reboul: Software (ClearSMILES development and analysis code), Methodology (ClearSMILES), Investigation, Writing - Review and Editing Antoine Taly: Supervision, Writing - Review and Editing Jérôme Waldispühl: Supervision, Writing - Review and Editing

## 7 Supporting information

- Table 1: SELFIES symbol-to-index mapping for numerical tokens.
- Table 2: Token loss categorisation for 300K sample batches
- generated by each augmented SELFIES VAE.
- Table 3: Stability and Molecular Descriptors for Different Models.
- Table 4: Deciles of Number of Equivalent ClearSMILES per original SMILES.
- Table 5: Proportion of Delta Values for the number of branches between corresponding sampled ClearSMILES and canonical SMILES.
- Table 6: Tokens per string quantiles for sampled ClearSMILES, RDKit default canonical SMILES, and canonical Kekulé SMILES.
- Table 7: Demi-decile of the branch size for ClearSMILES and SMILES.
- Table 8: SMILES fidelity analyses.
- Table 9: Viabilitiy metric (novelty, uniqueness, Validity) for the 300k samples generated by canonical SMILES VAE with 15 latent dimensions.
- Table 10: Viabilitiy metric (novelty, uniqueness, Validity/stability) for the 300k samples generated by SELFIES based VAE with 15 latent dimensions.
- Figure 1: kernel density estimation of the token loss for each 300 k samples batches generated from SELFIES-based VAEs.
- Figure 2: kernel density estimation of the token loss for each 300 k samples batches generated from MolGPT.
- Figure 3: Countplot of mutation type for each models.
- Figure 4: Heatmap of the absolute change between rings features normalized contingency table of MOSES and 300K samples generated from SELFIES-based VAE with 22 latent dimensions.
- Figure 5: Heatmap of the absolute change between rings features normalized contingency table of MOSES and 300K samples generated from SELFIES-based VAE with 15 latent dimensions.
- Figure 6: Heatmap of the absolute change between rings features normalized contingency table of MOSES and 300K samples generated from selfies GPT models with sampling temperature of 1.0.
- Figure 7: Heatmap of the absolute change between rings features normalized contingency table of MOSES and 300K samples generated from selfies GPT models with sampling temperature of 1.5.
- Figure 8: Histogram of number of tokens per string for SMILES.
- Figure 9: Duration distribution for processing a SMILES into a ClearSMILES.
- Figure 10: Contingency table of MOSES ring feature presented as Heatmap.
- Figure 11: Heatmap of the absolute change between rings features normalized contingency table of MOSES and 300K samples generated for SMILES representation by VAE with 22 latent dimensions.
- Figure 12: Heatmap of the absolute change between rings features normalized contingency table of MOSES and 300K samples generated for SMILES representation by VAE with 15 latent dimensions.
- Figure 13: Heatmap of the absolute change between rings features normalized contingency table of MOSES and 300K samples generated for SMILES representation by MolGPT with a sampling temperature of 1.0.
- Figure 14: Heatmap of the absolute change between rings features normalized contingency table of MOSES and 300K samples generated for SMILES representation by MolGPT with a sampling temperature of 1.5.
- Figure 15: Shannon Information Entropy for each SMILES based models across 100 epochs of training.
- Figure 16: Error of samples generated with MolGPT using a sampling temperature of 1.5 for SMILES.
- Figure 17: Error of samples generated with MolGPT using a sampling temperature o 1.0 for SMILES.
- Figure 18: Error of samples generated with VAE with 15 latent dimensions for SMILES.
- Figure 19: Assessment of various fidelity metrics for all viable samples generated by all VAE models with 15 latent dimensions.
- Figure 20: Assessment of various fidelity metrics for all viable samples generated by MolGPTs models with sampling temperature of 1.0.

## 10 Data and Software Availability

The code for the ClearSMILES pipeline can be found in github: https://github.com/EtienneReboul/ClearSMILES

The training data, models checkpoints and samples can be found on zenodo: https://doi.org/10.5281/zenodo.14420504 The SELFIES string stability testing is available as a jupyter notebook on google colab at: https://colab.research.google.com/drive/1jZwRGyXqUSaQhQ-yRILBoqiM1HM1ikK0?usp=sharing

