## Supplementary Information for "Improving the reliability of molecular string representations for generative chemistry"

### SI :SELFIES

#### additional documentation

SELFIES grammar uses overloading to limit the number of unique tokens needed to encode the length of branches and rings as hexadecimal. The following table 1 describes the mapping between the overloaded token and their hexadecimal equivalent.

The original SELFIES have a vocabulary composed of 23 tokens, the same size as the vocabulary of canonical SMILES. The SELFIES vocabulary contains the following tokens : '[#Branch1]', '[#Branch2]', '[#C]','[#N]', '[=Branch1]', '[=Branch2]', '[=C]', '[=N]', '[=O]', '[=Ring1]', '[=Ring2]', '[=S]', '[Br]', '[Branch1]', '[Branch2]', '[C]', '[Cl]', '[F]', '[NH1]', '[N]','[O]', '[P]', '[Ring1]', '[Ring2]', '[S]'

For the vocabulary of SELFIES without overloading, the token "[P]" was removed as MOSES does not contain molecules with Phosphorus and was only used for indexing; 16 non-overloaded tokens were added to the vocabulary ([0]...[15]). The "[P]" token was also

Table 1: symbol-to-index mapping for numerical tokens, this table was taken from SELFIES documentation

| Index | Symbol | Index | Symbol |
|---|---|---|---|
| 0 | [C] | 8 | [#Branch2] |
| 1 | [Ring1] | 9 | [O] |
| 2 | [Ring2] | 10 | [N] |
| 3 | [Branch1] | 11 | [=N] |
| 4 | [=Branch1] | 12 | [=C] |
| 5 | [#Branch1] | 13 | [#C] |
| 6 | [Branch2] | 14 | [S] |
| 7 | [=Branch2] | 15 | [P] |

All other symbols are assigned index 0

removed from the SELFIES vocabulary, with the additional regular branch and ring tokens: [#Branch1]', [#Branch2]', [=Branch1]', [=Branch2]', [=Ring1]', [=Ring2]', [=Ring1]', [=Ring2]', [Branch1]', [[Ring2]', [[Ring1]', [Ring2]'. In total, 11 tokens were removed from the original SELFIES vocabulary. Compared to SELFIES without overloading, an additional 24 explicit numerical tokens were introduced into the SELFIES without hexadecimal vocabulary (16-39). The only explicit numerical token from the SELFIES with no overloading not used by the hexadecimal SELFIES is "[0]". An additional 4 non numbered branch and ring tokens were added to the vocabulary: [#Branch]', '[=Branch]', '[=Ring]', '[Branch]', '[Ring]'.

#### SELFIES error analysis

We measured the extent of information loss during the decoding process of the SELFIES algorithm. To that end, we quantified the token loss between the sampled SELFIES and the regenerated SELFIES. The kernel density estimation of the token loss for each SELFIES-based VAE is illustrated in Figure 1. Our findings reveal that the SELFIES samples exhibit a similar multimodal distribution with minor variations among the different models.

We found that 92% of the unstable samples generated by the SELFIES-based models had a token loss below zero, which means that there was less token in the regenerated

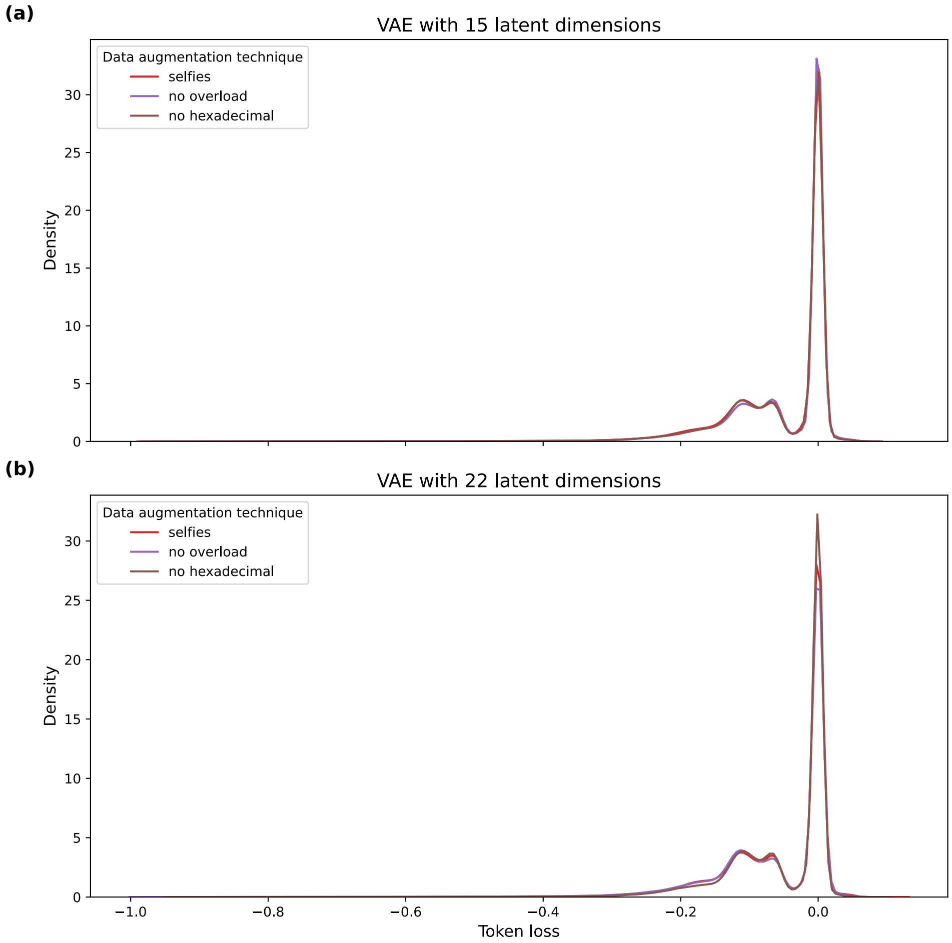


Figure 1: kernel density estimation of the token loss for each 300 k samples batches generated from SELFIES-based VAEs

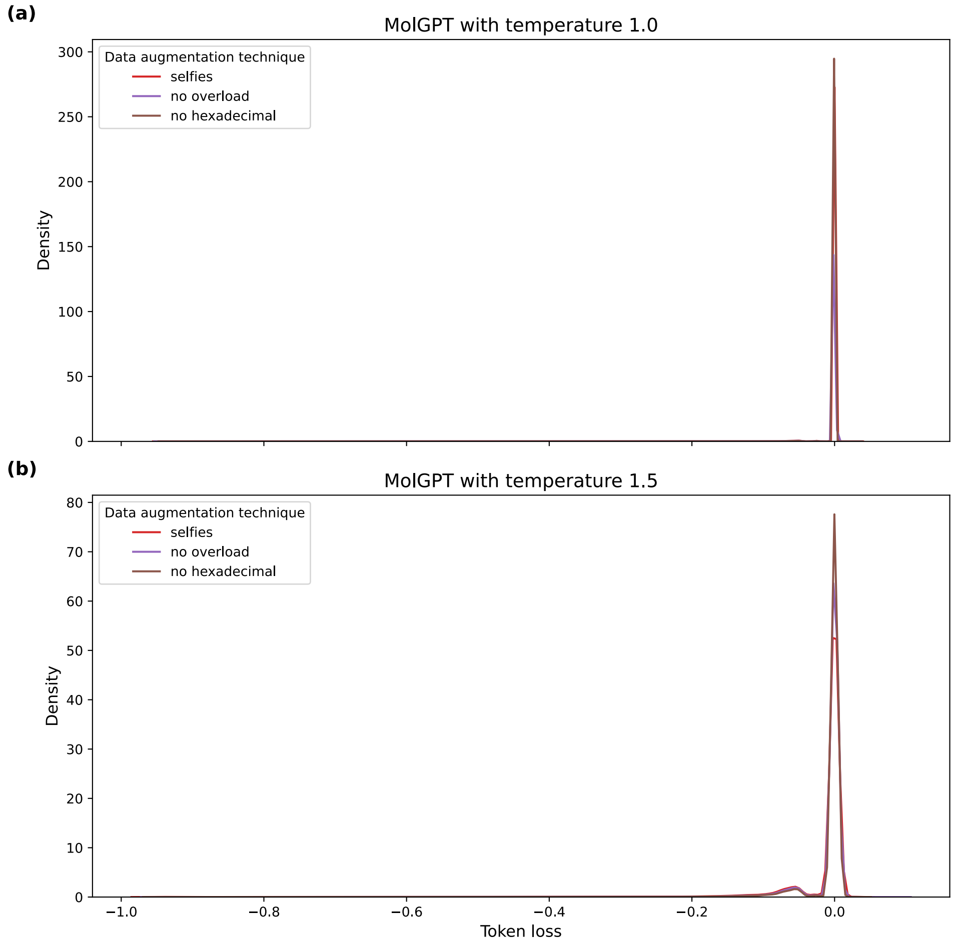


Figure 2: kernel density estimation of the token loss for each 300 k samples batches generated from MolGPT

SELFIES than in the original samples. The details of augmented SELFIES are described in the Supplementary Table 2. This suggests that in an overwhelming majority of cases, unstable SELFIES exhibit a loss of information during their translation to SMILES. We found that in 4. 5% of unstable SELFIES there was a token loss equal to zero, meaning that the number of tokens was not modified between regenerated SELFIES and the original samples. This could indicate that an overloaded token was swapped for another or that the number of bonds was adjusted for an atom token , such as a double bond token ([=C]) was adjusted to a simple bond token ([C]). Interestingly, in 3. 5% of the cases, there was a token loss superior to zero, meaning an actual token gain between regenerated SELFIES and the original samples. This might be linked to a renumbering where more hexadecimal tokens are needed to describe either a branch or a ring closure.

Table 2: Token loss categorisation for 300K sample batches generated by each augmented SELFIES VAE

| **Data augmentation** | **Latent Dimensions** | **Loss Label** | **Proportion (%)** |
|---|---|---|---|
| `No hexadecimal` | 15 | `loss` | 92.0 |
| | | `gain` | 4.2 |
| | | `unchanged` | 3.8 |
| `No hexadecimal` | 22 | `loss` | 93.3 |
| | | `gain` | 3.6 |
| | | `unchanged` | 3.1 |
| `No overload` | 15 | `loss` | 90.5 |
| | | `gain` | 5.4 |
| | | `unchanged` | 4.1 |
| `No Overload` | 22 | `loss` | 92.0 |
| | | `gain` | 4.3 |
| | | `unchanged` | 3.7 |

We made alignment between the original samples and their regenerated counterpart with the Needleman-Wunsh algorithm (string2string Python module) to further our understanding of the SELFIES instability. We classified mutations according to the classical pattern of deletion, substitution, and addition. The count of mutations per SELFIES is available in the supplementary figure 3. The most prevalent type of mutation is deletion, followed by substitution. Insertion mutation are comparatively rare compared to deletion and substitution. An important thing to note is that the mutations are not mutually exclusive. Therefore, one or more deletions, insertions, or substitutions can occur simultaneously. The counts of mutations reveals that the most common scenario is one mutation per type per SELFIES samples. The number of mutations of the same type is inversely proportional to its occurrence. Having 3 or more mutations of the same type per SELFIES is rare for deletion and substitution. We did not observe more than 2 insertions per SELFIES, with 2 insertions in the same SELFIES being almost negligible.

To go further into the analysis of mutation, we annotated the token into four different categories : "a" for atom tokens ([C]) , "b" for branch token ([Branch1]) , "r" for ring token ([=Ring1]), "n" for numerical/overloaded tokens ([Branch1]**[C]**). The most common deletion was the loss of ring tokens and the associated overloaded tokens, i.e., ring closure. Closely following is the loss of branches tokens associated with their overloaded or atom sequences depending on the number of latent dimensions used in the model. The number of deletions for the two deletion types is of the same order of magnitude in samples generated by SELFIES based VAE(s) with 15 or 22 latent dimensions. This result suggests that VAE struggles to understand the syntax of SELFIES for both ring closure and branches.

To confirm this hypothesis, we performed a more in-depth analysis of mutations based on token classification, categorizing tokens as atoms, numerical, branches, overloaded numerical token, or rings. The detailed classification is available in a CSV file in the supplementary data. Our analysis revealed that the most commonly deleted tokens are those composed of a ring token and one or more overloaded tokens, indicating the deletion of ring closure in the molecules. This trend is consistent across all data-augmentation procedures, highlighting the significant challenge in properly representing rings in SELFIES for our VAEs models. Additionally, the most frequently deleted tokens include branches tokens associated with one or more overloaded tokens, branches tokens alone, and atoms tokens. The order of

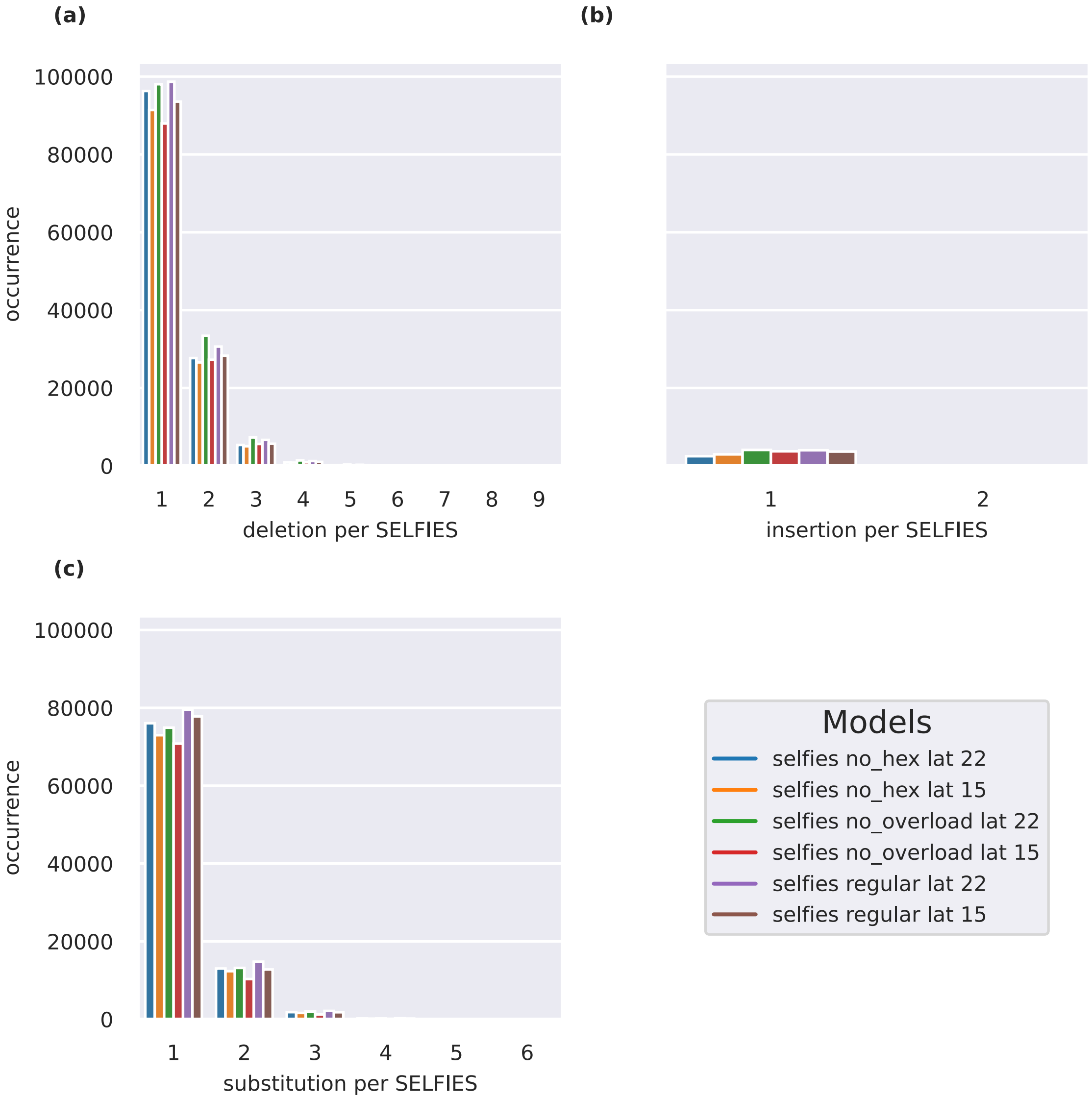


Figure 3: Countplot of mutation type for each models, (a) number of Deletions per SELFIES , (b) number of insertion per SLEFIES (c) number of Substitution per SELFIES

the most prevalent mutation type changes depending on the data augmentation procedure, but we could not find a significant pattern. In the case of the deletion of branches with overloaded tokens, it implies that the atomic content of these branches is reallocated to the main chain, essentially inserting the atoms from the faulty branch into the main chain. Regarding the deletion of branches tokens only, it signifies that one or multiple overloaded tokens following the branch token are de-overloaded and interpreted as atoms, also being reallocated to the main chain.Finally, the deletion of atoms tokens occurs because they do not adhere to the theoretical valence limits. This indicates that VAEs still face challenges in maintaining consistent valence throughout the entire molecules when dealing with SELFIES representations.

#### SELFIES fidelity analysis

As the most deleted feature was the ring closure, we performed the same analysis based on the contingency table of the ring features as with the canonical SMILES. The results of this analysis are presented in Supplementary Figure 4 (a) and in Supplementary Figure 5 (a) for samples generated by SELFIES based VAE with 22 or 15 latent dimensions, respectively. The results of the ring feature analysis show that the issues found in canonical SMILES samples persist but are also aggravated. The drop in ring with 6 atoms in a molecule with 3 or more rings is 3 times the drop with canonical SMILES, to reach a 10.8 percentage point difference with the MOSES baseline.

The outliers rings are more frequent in samples generated by SELFIES VAE with 22 or 15 latent dimensions, as they represent around 7.5% rings. The degree of "aberrancy" of rings is also starker in SELFIES samples with as many as 22 atoms per rings, whereas canonical SMILES have a maximum ring size of 13 atoms. The size of the outlier ring is also inversely proportional to its occurrence, suggesting that the more aberrant a ring or molecule is , the less frequent it is.

To quantify the functional impact of the instability of SELFIES on the fidelity of the

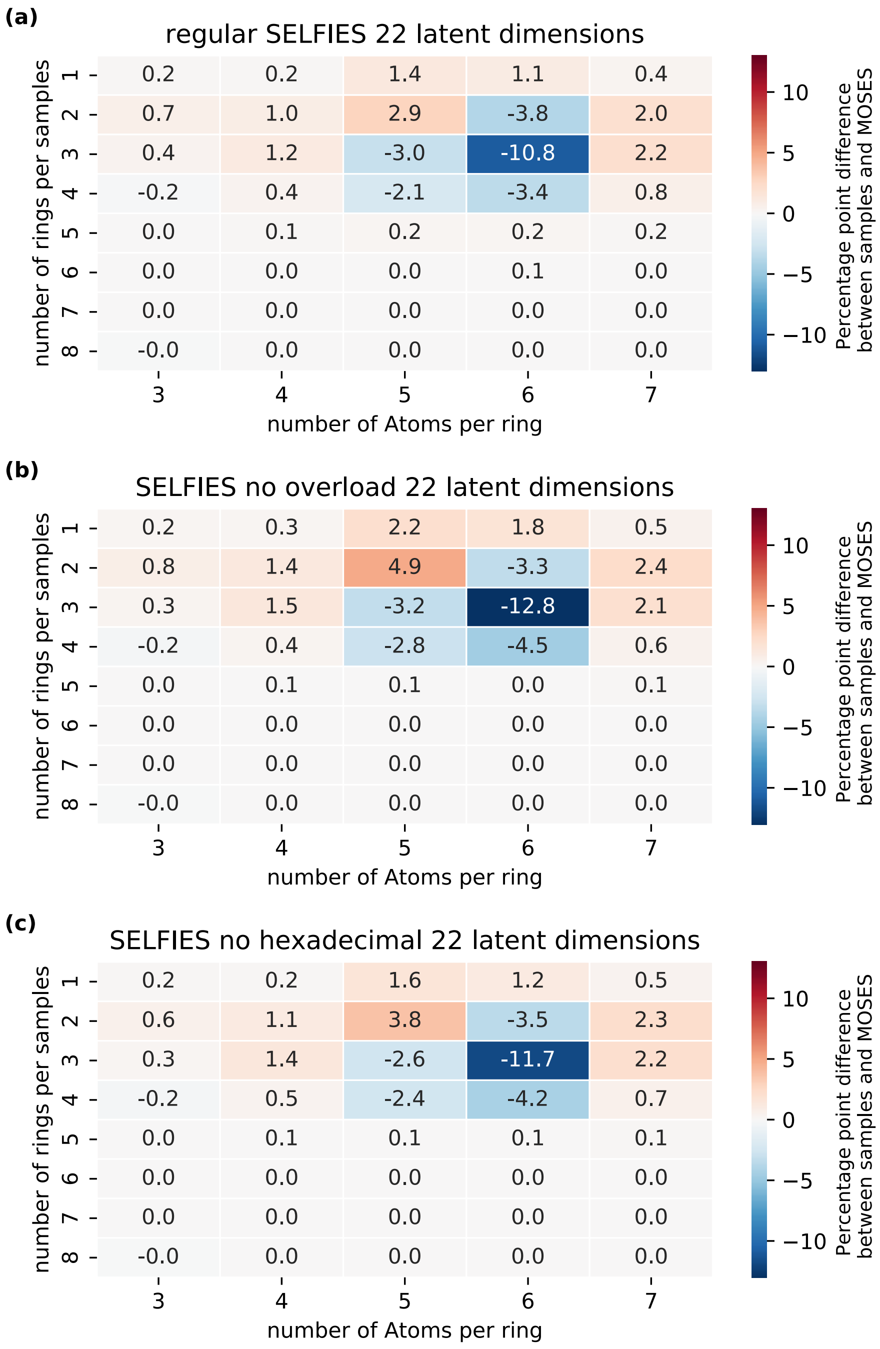


Figure 4: Heatmap of the absolute change between rings features normalized contingency table of MOSES and 300K samples generated from (a) SELFIES-based VAE (b) SELFIES with no overload based VAE (c) SELFIES with no hexadecimal numbering, both with 22 latent dimensions.

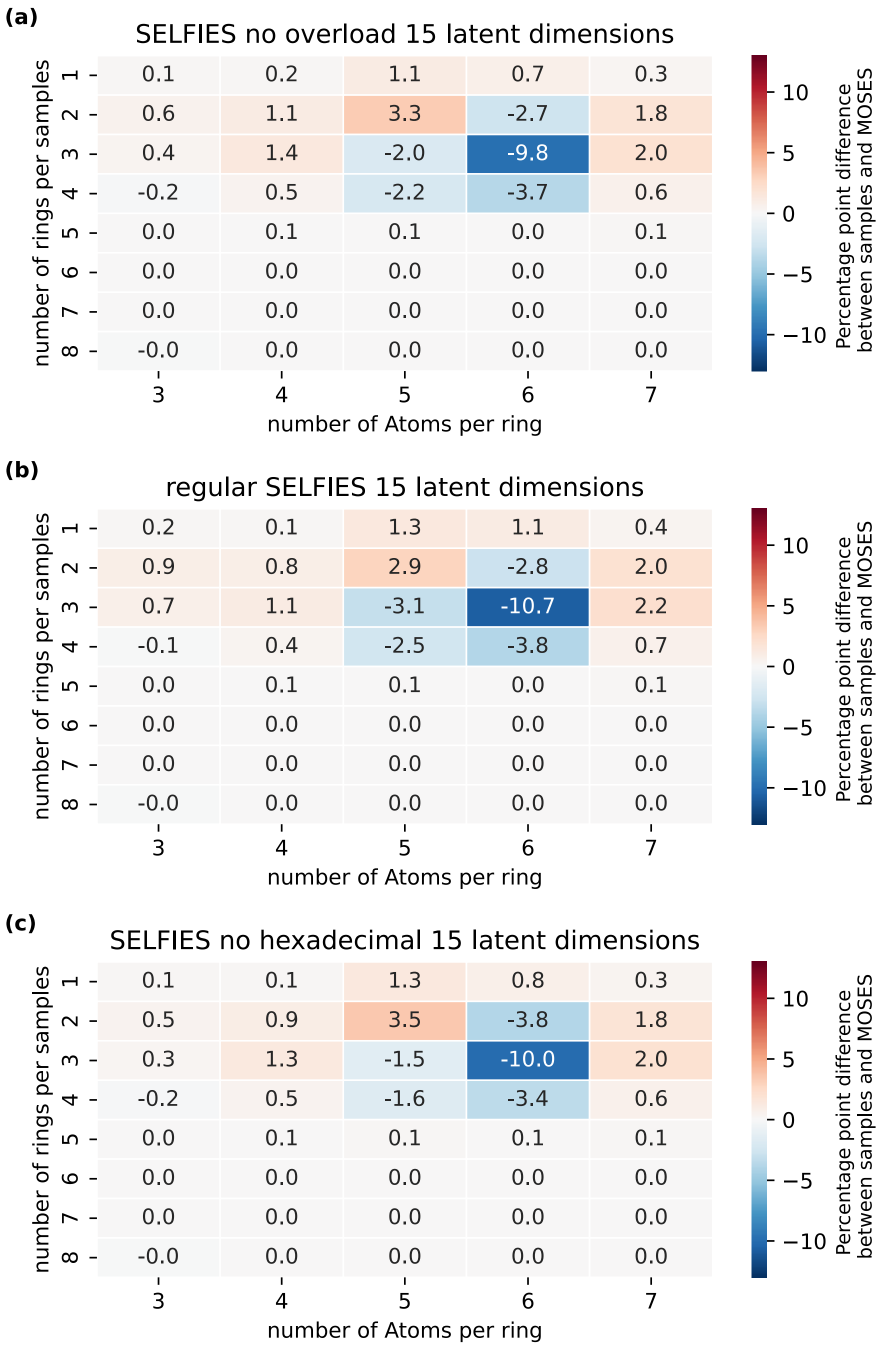


Figure 5: Heatmap of the absolute change between rings features normalized contingency table of MOSES and 300K samples generated from (a) SELFIES-based VAE (b) SELFIES with no overload based VAE (c) SELFIES with no hexadecimal numbering, both with 15 latent dimensions.

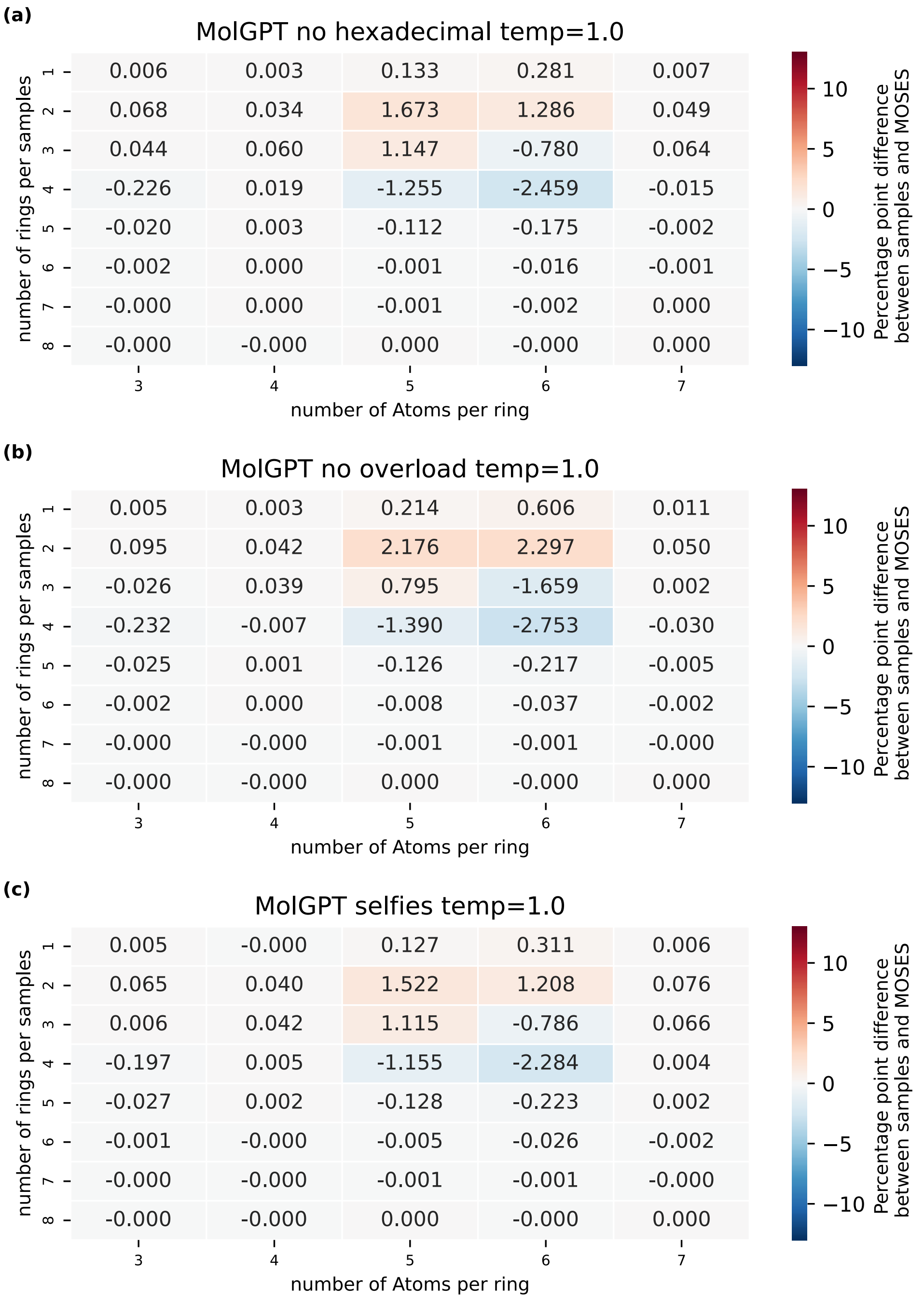


Figure 6: Heatmap of the absolute change between rings features normalized contingency table of MOSES and 300K samples generated from selfies GPT models with sampling temperature of 1.0

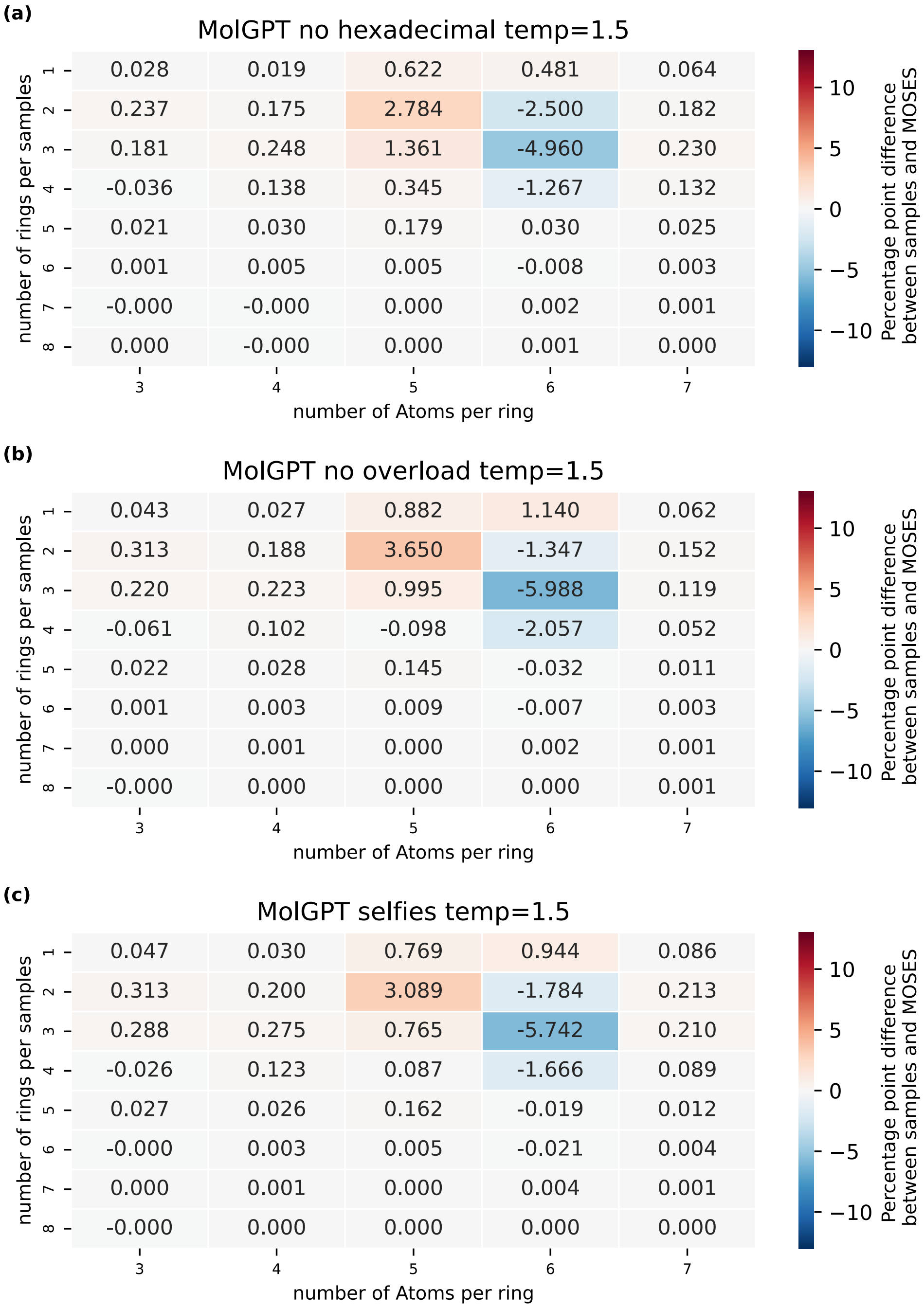


Figure 7: Heatmap of the absolute change between rings features normalized contingency table of MOSES and 300K samples generated from selfies GPT models with sampling temperature of 1.5

samples, we split the stable and unstable SELFIES into two different sets. For each, we computed the watertein distance for Molecular Weight, QED, SA, and TPSA. We found that instability leads to an overall increase in the wassertein distance, meaning that the loss or alteration caused by SELFIES effectively translates into a decrease in fidelity, as shown in Table 3:

Table 3: Stability and Molecular Descriptors for Different Models

| **Model** | **Stability** | **MolWeight** | **QED** | **SA** | **TPSA** |
|---|---|---|---|---|---|
| selfies no hex lat 22 | Stable | 7.269829 | 0.056104 | 1.109658 | 6.007697 |
| selfies no hex lat 22 | Unstable | 6.610271 | 0.170771 | 1.514347 | 9.940050 |
| selfies no hex lat 15 | Stable | 5.520926 | 0.053395 | 0.956029 | 5.917653 |
| selfies no hex lat 15 | Unstable | 6.837464 | 0.169941 | 1.400146 | 10.977418 |
| selfies no overload lat 22 | Stable | 6.878443 | 0.072431 | 1.093295 | 7.673425 |
| selfies no overload lat 22 | Unstable | 8.403700 | 0.199772 | 1.507760 | 12.335939 |
| selfies no overload lat 15 | Stable | 4.560730 | 0.056527 | 0.969116 | 5.006249 |
| selfies no overload lat 15 | Unstable | 6.285555 | 0.167162 | 1.371182 | 7.914755 |
| selfies regular lat 22 | Stable | 6.447442 | 0.052974 | 1.056126 | 4.163644 |
| selfies regular lat 22 | Unstable | 7.532643 | 0.160952 | 1.426262 | 4.814064 |
| selfies regular lat 15 | Stable | 5.930321 | 0.056284 | 1.007180 | 3.699631 |
| selfies regular lat 15 | Unstable | 8.476067 | 0.158868 | 1.440207 | 4.702175 |

### SI :SMILES

#### Vocabulary of canonical SMILES

The vocabulary of canonical SMILES generated by the RDKIT module consists of 23 tokens: #, ’(’,’)’,’-,’1’,’2’,’3’,’4’,’5’,’6’,’=’,’Br’,’C’,’Cl’,’F’,’N’,’O’,’S’,’[nH]’,’c’,’n, o’, s’. The original canonical SMILES of the MOSES dataset has an extra token: [H]’.

#### additional figures on ClearSMILES properties

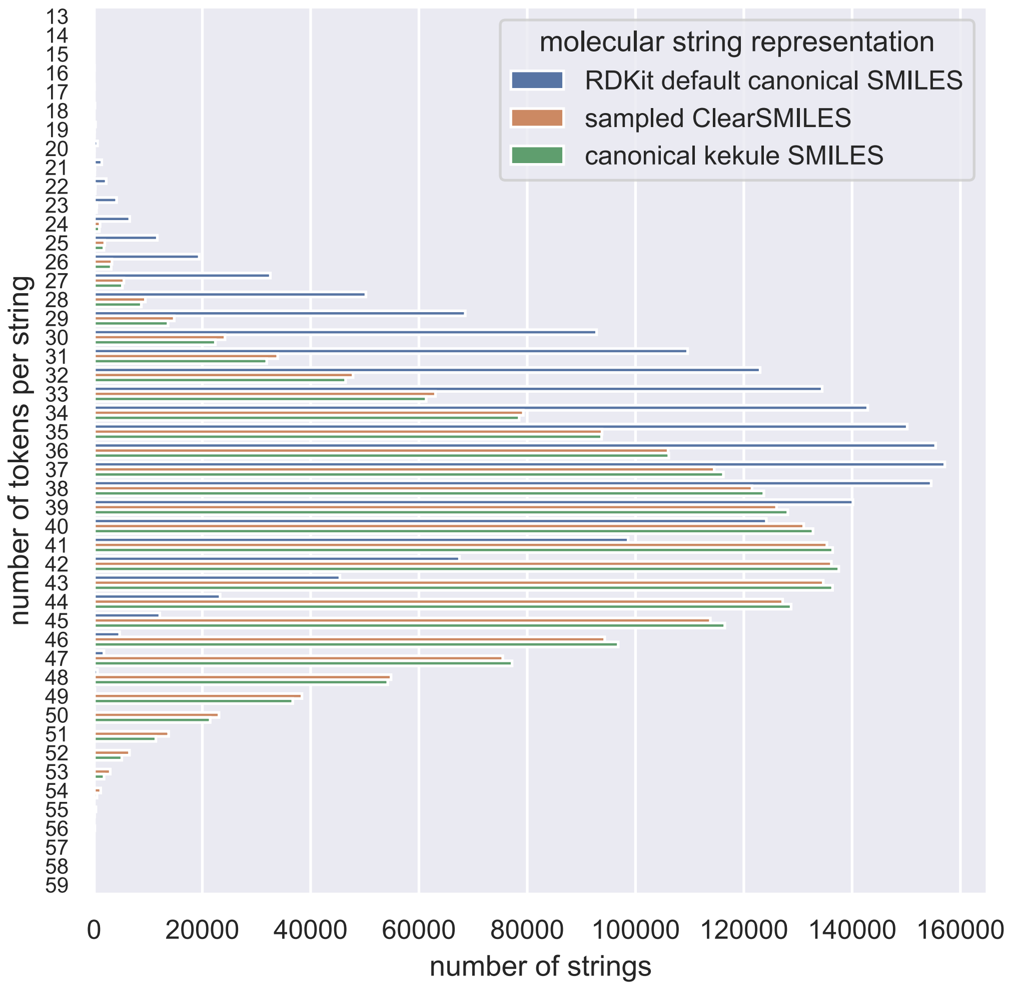


Figure 8: Histogram of number of tokens per string for RDKit default canonical SMILES (blue) , ClearSMILES (orange), canonical kekule SMILES (green)

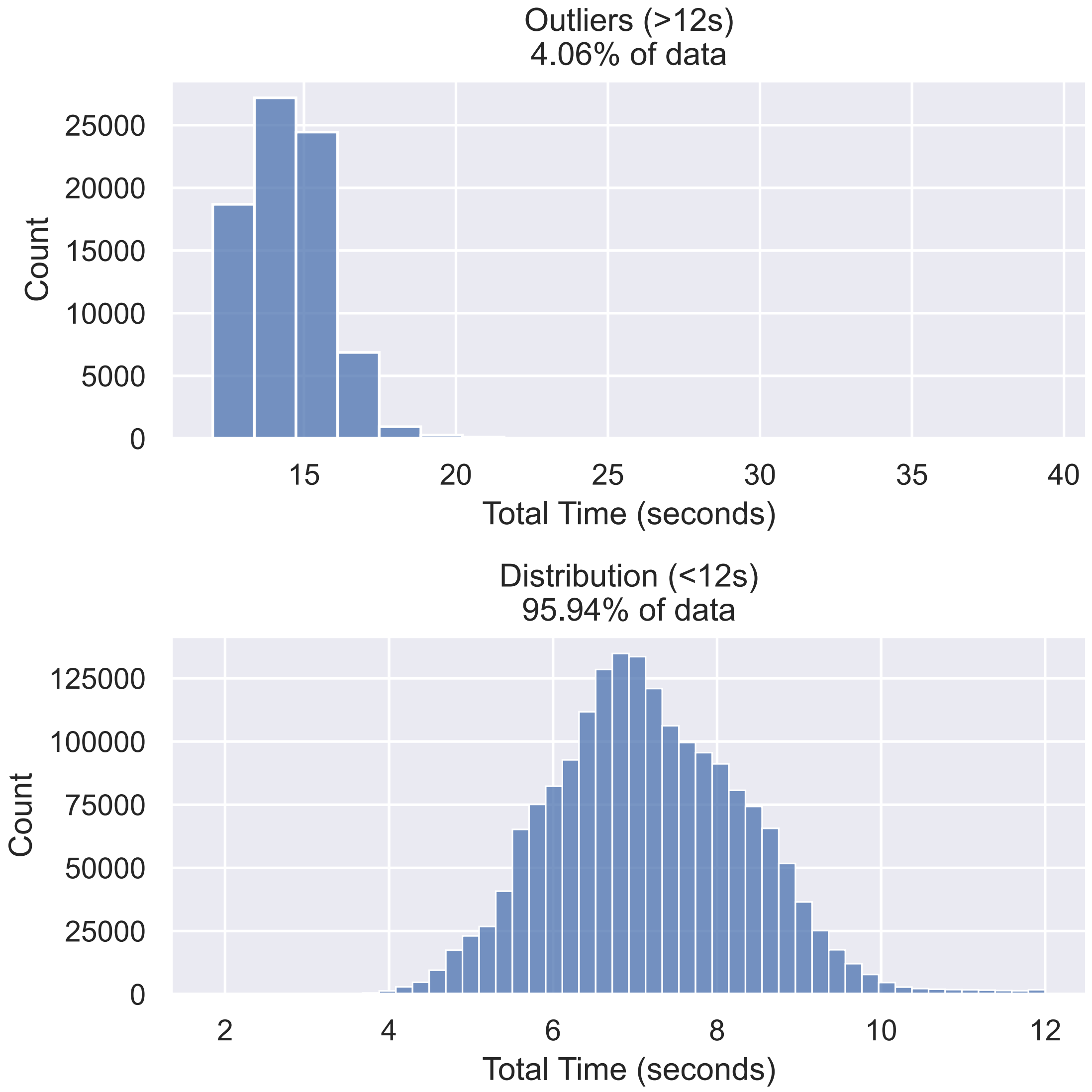


Figure 9: Duration distribution for processing a SMILES into a ClearSMILES. The top panel shows the distribution of outliers, which account for approximately 4% of the data. The bottom panel presents the distribution for the main group, representing about 95% of the data.

Table 4: Deciles of Number of Equivalent ClearSMILES per original SMILES

| Decile | Number of Equivalent Solutions |
|---|---|
| 0.0 | 1 |
| 0.1 | 1 |
| 0.2 | 2 |
| 0.3 | 2 |
| 0.4 | 2 |
| 0.5 | 2 |
| 0.6 | 4 |
| 0.7 | 4 |
| 0.8 | 4 |
| 0.9 | 8 |
| 1.0 | 128 |

Table 5: Proportion of Delta Values for the number of branches between corresponding sampled ClearSMILES and canonical SMILES

| Δ Branches | Proportion (%) |
|---|---|
| -3 | 3.36e-3% |
| -2 | 4.59e-1% |
| -1 | 11.72% |
| 0 | 74.90% |
| 1 | 11.69% |
| 2 | 1.13% |
| 3 | 3.95e-2% |
| 4 | 6.20e-4% |

Table 6: Tokens per string quantiles for sampled ClearSMILES, RDKit default canonical SMILES, and canonical Kekulé SMILES

| Quantile | RDKit Canonical SMILES | Canonical Kekulé SMILES | Sampled ClearSMILES |
|---|---|---|---|
| 0.00 | 13.0 | 13.0 | 13.0 |
| 0.25 | 32.0 | 37.0 | 36.0 |
| 0.50 | 36.0 | 40.0 | 40.0 |
| 0.75 | 39.0 | 44.0 | 44.0 |
| 1.00 | 54.0 | 57.0 | 59.0 |

Table 7: demi-decile of the branch size for ClearSMILES and SMILES

| Quantile | ClearSMILES | SMILES |
|---|---|---|
| 0.00 | 0.0 | 0.0 |
| 0.05 | 1.0 | 2.0 |
| 0.10 | 2.0 | 2.0 |
| 0.15 | 2.0 | 2.0 |
| 0.20 | 2.0 | 2.0 |
| 0.25 | 2.0 | 3.0 |
| 0.30 | 3.0 | 3.0 |
| 0.35 | 3.0 | 3.0 |
| 0.40 | 3.0 | 3.0 |
| 0.45 | 3.0 | 3.0 |
| 0.50 | 3.0 | 3.0 |
| 0.55 | 3.0 | 4.0 |
| 0.60 | 4.0 | 4.0 |
| 0.65 | 4.0 | 4.0 |
| 0.70 | 4.0 | 4.0 |
| 0.75 | 4.0 | 4.0 |
| 0.80 | 4.0 | 4.0 |
| 0.85 | 5.0 | 5.0 |
| 0.90 | 5.0 | 5.0 |
| 0.95 | 6.0 | 6.0 |
| 1.00 | 11.0 | 11.0 |

#### additional figures on ring feature properties

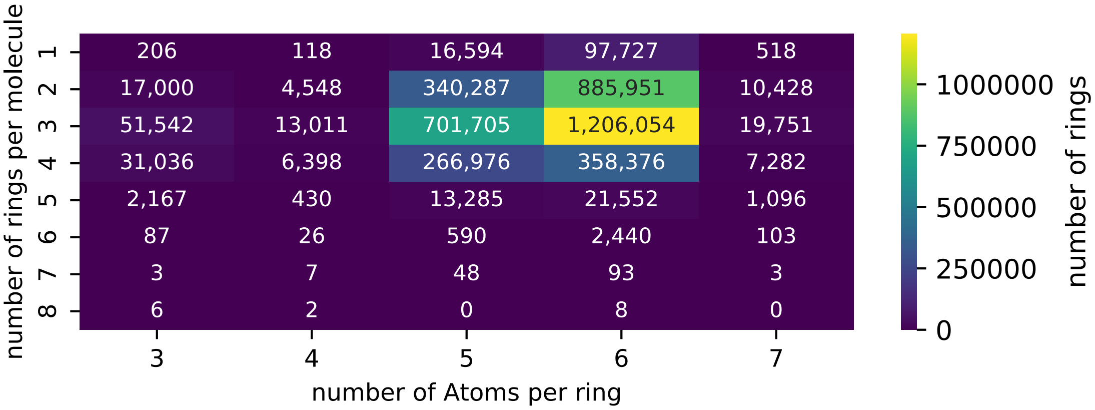


Figure 10: Contingency table of MOSES ring feature presented as Heatmap. The ring feature are the number of rings per molecule (rows) and the number of atoms per ring (columns).

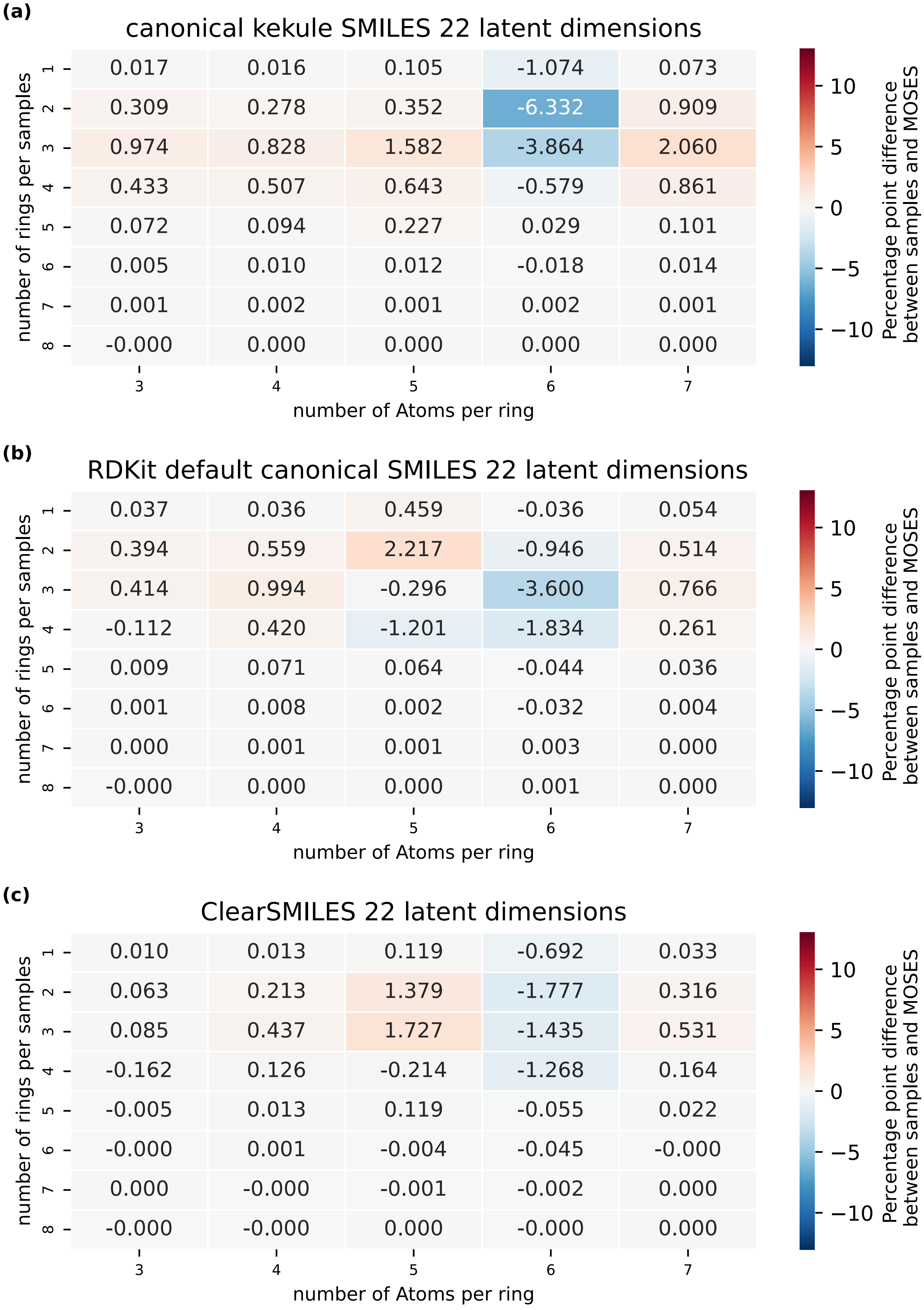


Figure 11: Heatmap of the absolute change between rings features normalized contingency table of MOSES and 300K samples generated for clearsmiles , RDKit default canonical smiles and canonical kekule SMILES representation by VAE with 22 latent dimensions.

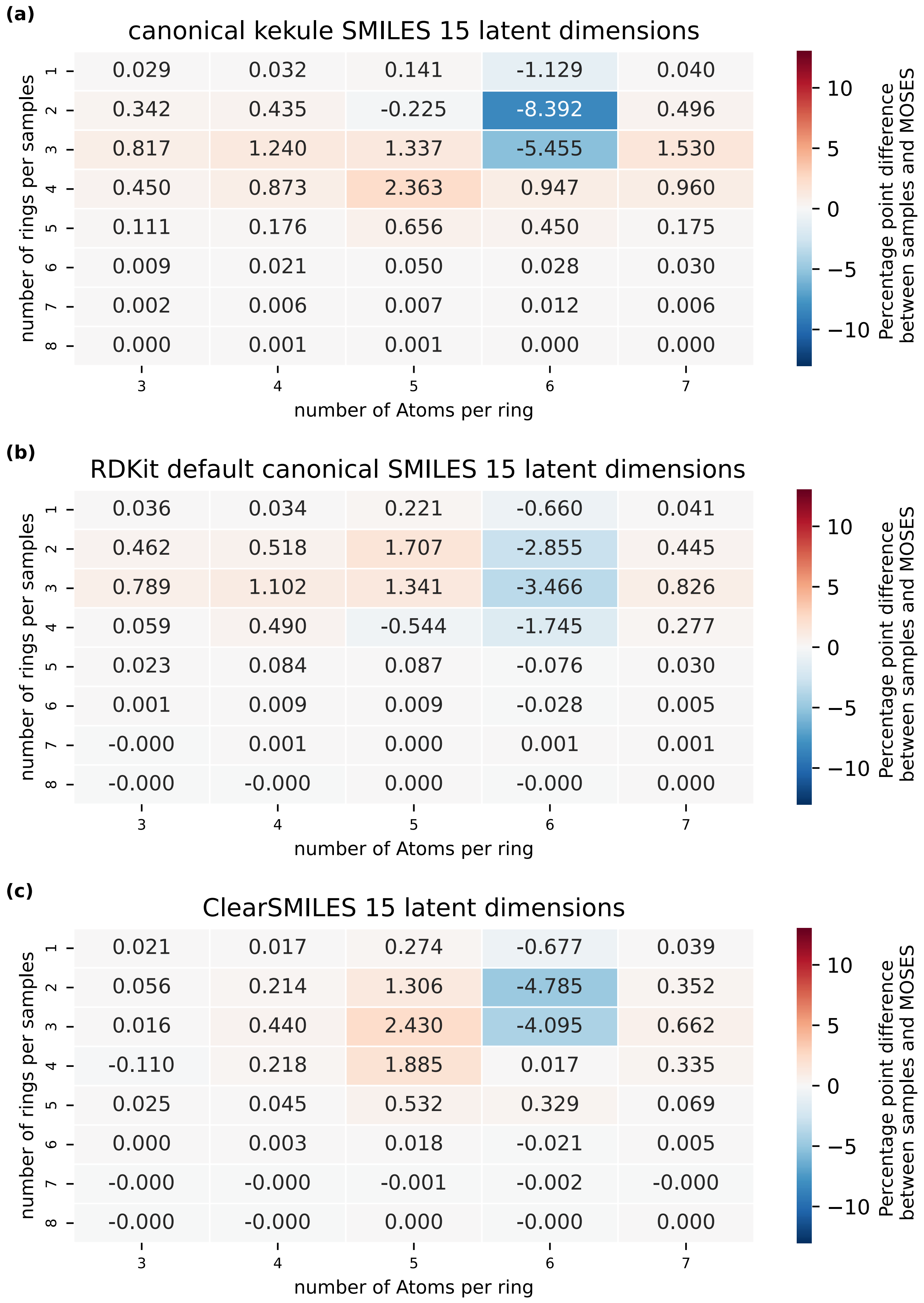


Figure 12: Heatmap of the absolute change between rings features normalized contingency table of MOSES and 300K samples generated for clearsmiles , RDKit default canonical smiles and canonical kekule SMILES representation by VAE with 15 latent dimensions.

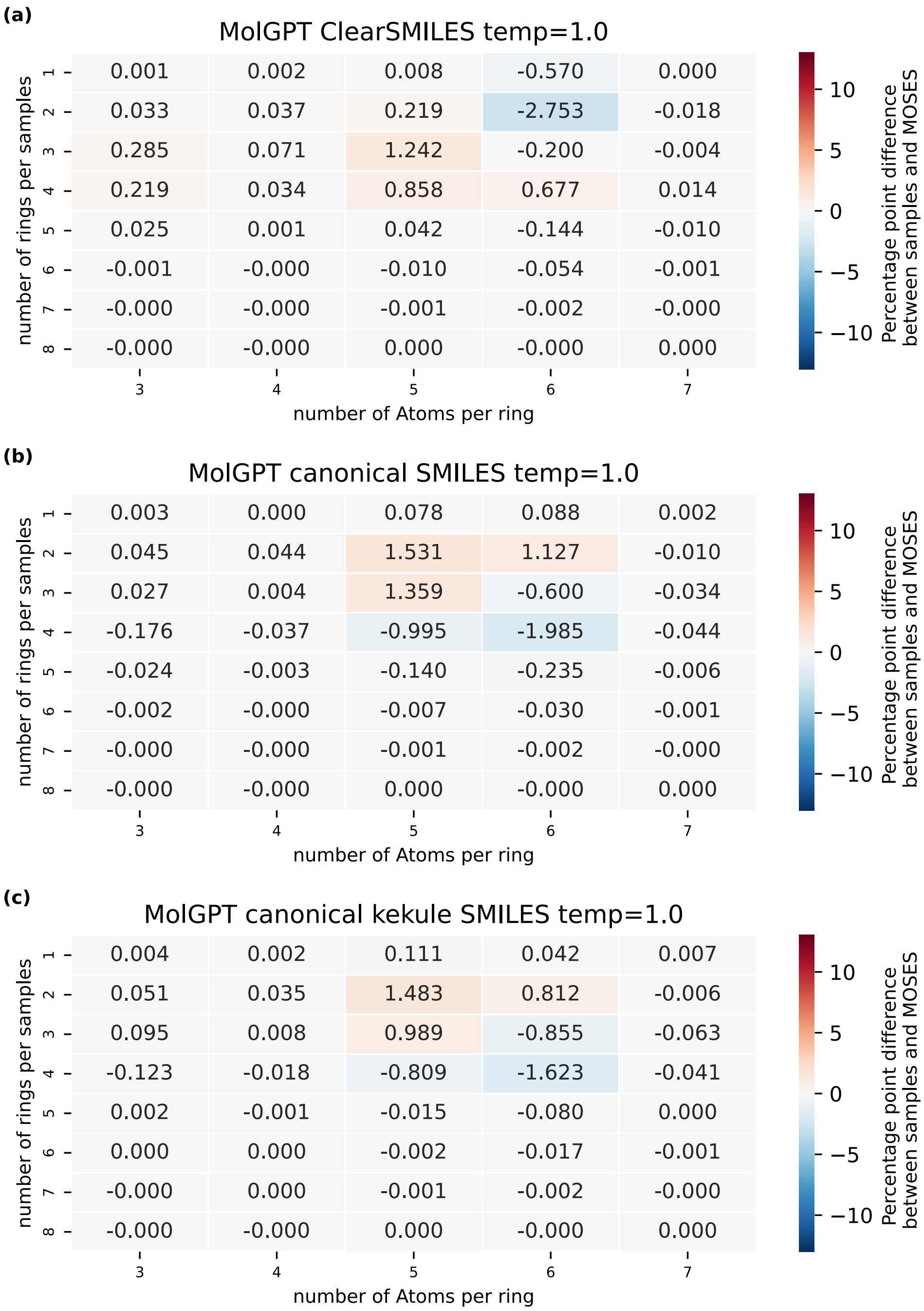


Figure 13: Heatmap of the absolute change between rings features normalized contingency table of MOSES and 300K samples generated for clearsmiles , RDKit default canonical smiles and canonical kekule SMILES representation by MolGPT with a sampling temperature of 1.0.

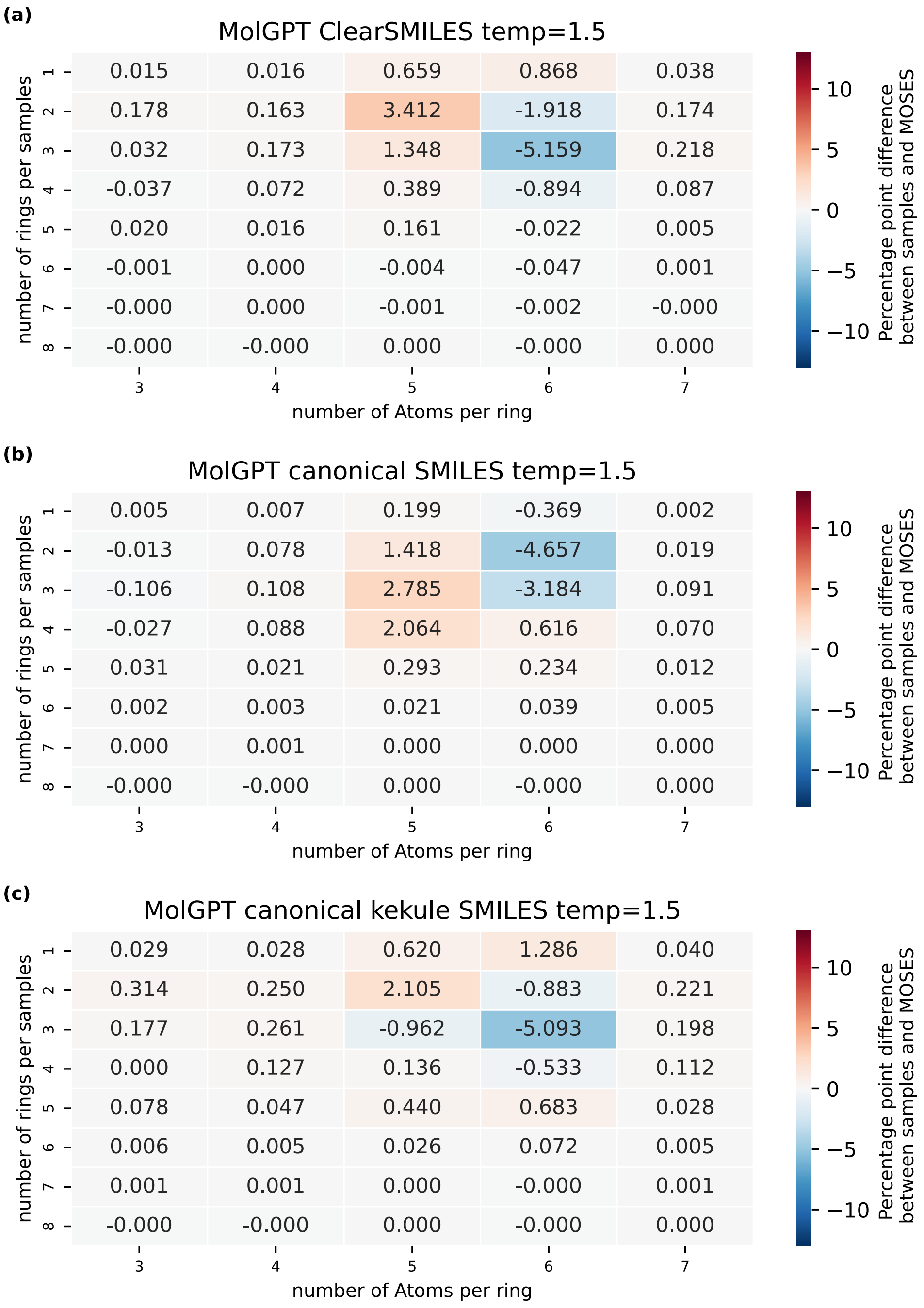


Figure 14: Heatmap of the absolute change between rings features normalized contingency table of MOSES and 300K samples generated for clearsmiles , RDKit default canonical smiles and canonical kekule SMILES representation by MolGPT with a sampling temperature of 1.5.

#### SMILES fidelity analyses

Table 8

| Model | Latent Dimensions | TPSA | MolWeight | QED | SA |
|---|---|---|---|---|---|
| SELFIES no hexadecimal | 22 | 7.798 | 6.805 | 0.115 | 1.316 |
| SELFIES no hexadecimal | 15 | 8.314 | 5.342 | 0.111 | 1.175 |
| SELFIES no overload | 22 | 10.094 | 6.693 | 0.141 | 1.315 |
| SELFIES no overload | 15 | 6.243 | **5.259** | 0.108 | 1.158 |
| regular SELFIES | 22 | 4.363 | 7.017 | 0.112 | 1.258 |
| regular SELFIES | 15 | **4.136** | 7.245 | 0.109 | 1.231 |
| canonical SMILES | 22 | 4.716 | 7.149 | 0.041 | 0.430 |
| canonical SMILES | 15 | 4.631 | 6.154 | 0.040 | 0.525 |
| ClearSMILES | 22 | 4.304 | 7.344 | **0.022** | **0.345** |
| ClearSMILES | 15 | 5.612 | 7.075 | 0.045 | 0.494 |

### Miscellaneous

Table 9: viabilitiy metric novelty,uniqueness,Validity for the 300k samples generated by canonical SMILES a VAE with 15 latent dimensions. Every metrics is expressed in percentages

| augmentation | Validity | novelty | uniqueness |
|---|---|---|---|
| RDKit default canonical SMILES | 80.6 | 99.64 | 99.98 |
| ClearSMILES | 96.75 | 99.4 | 99.97 |
| canonical kekule SMILES | 93.33 | 99.73 | 99.88 |

Table 10: viabilitiy metric novelty,uniqueness,Validity/stability for the 300k samples generated by SELFIES based VAE with 15 latent dimensions. Every metrics is expressed in percentages

| Representation | augmentation | Stability | novelty | uniqueness |
|---|---|---|---|---|
| SELFIES | regular | 48.34 | 99.91 | 99.98 |
| SELFIES | no overload | 52.94 | 99.90 | 99.93 |
| SELFIES | no hexadecimal | 50.60 | 99.90 | 99.90 |

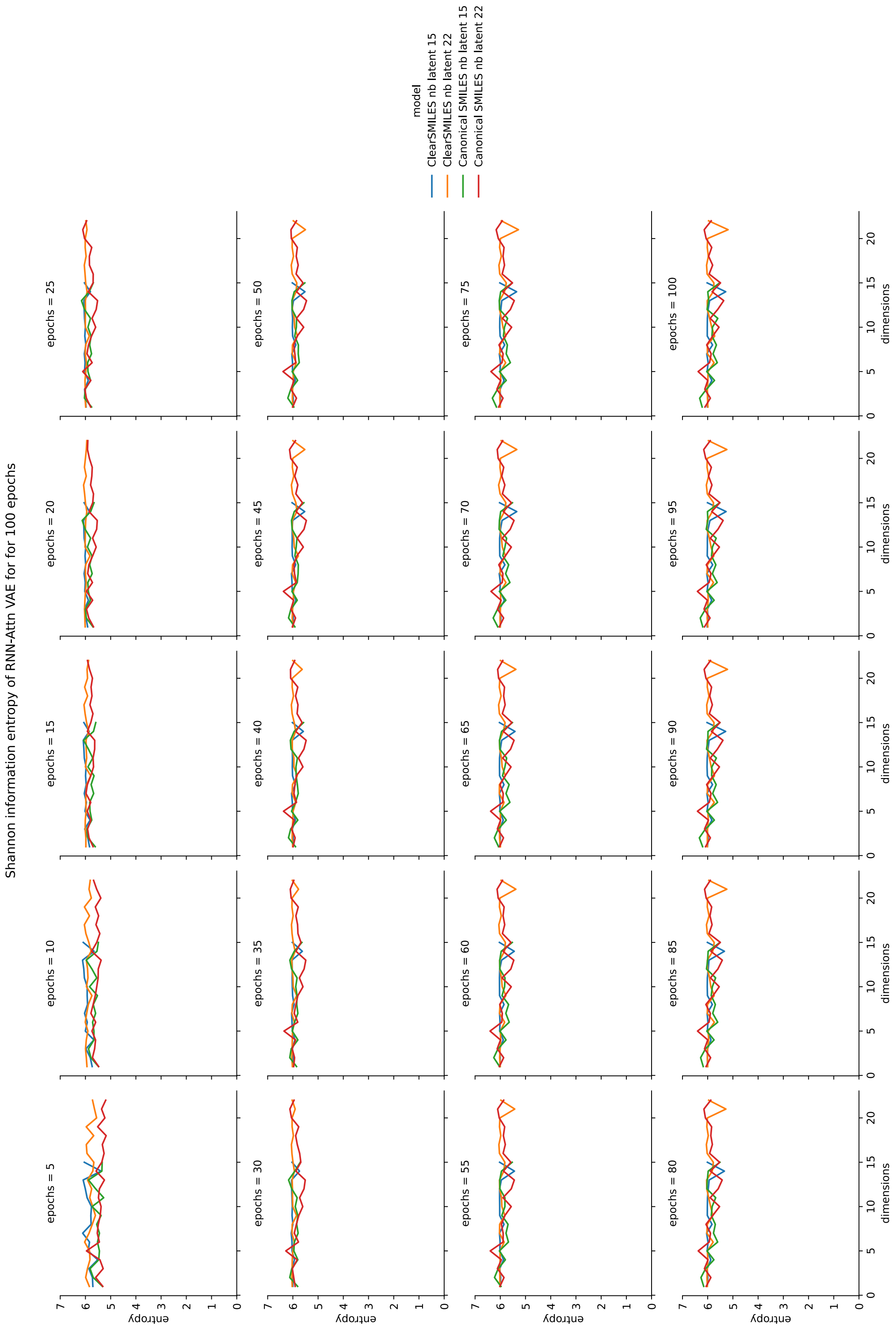


Figure 15: Shannon Information Entropy for each SMILES based models across 100 epochs of training

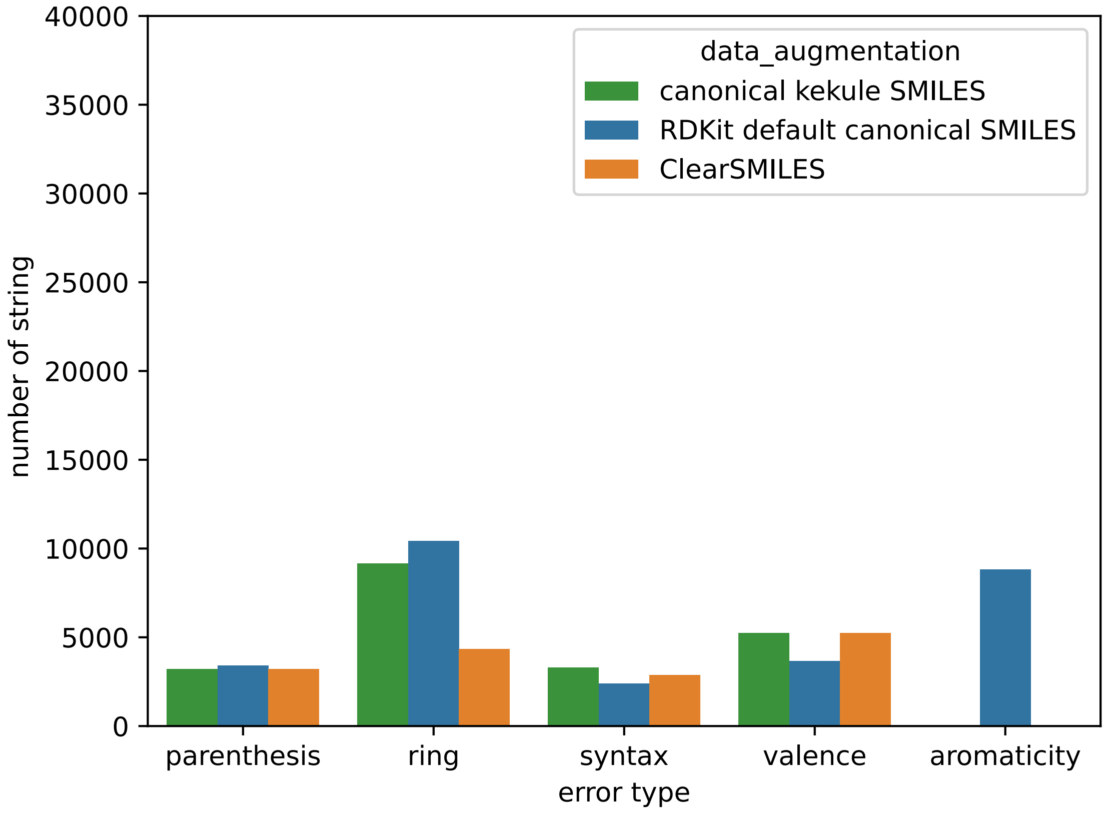


Figure 16: Error of samples generated with MolGPT using a sampling temperature of 1.5 for ClearSMILES (orange), RDKit Default canoncial SMILES, canonical kekule SMILES

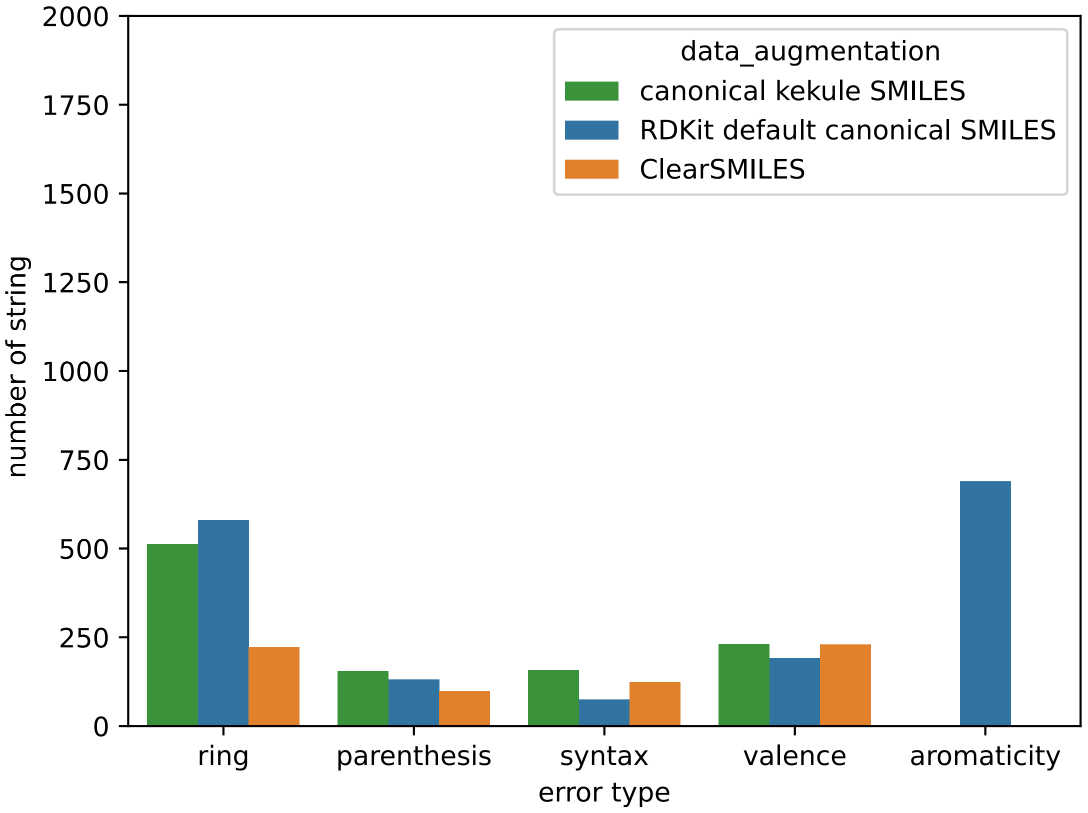


Figure 17: Error of samples generated with MolGPT using a sampling temperature of 1.0 for ClearSMILES (orange), RDKit Default canoncial SMILES, canonical kekule SMILES

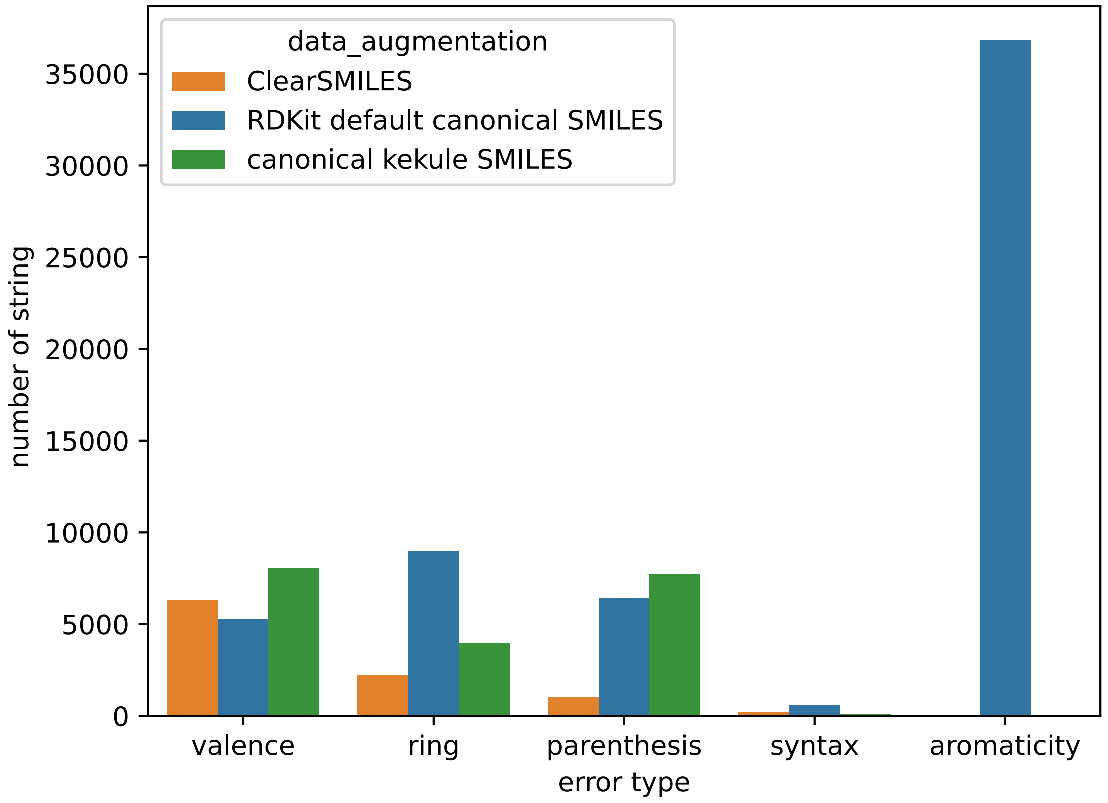


Figure 18: Error of samples generated with VAE with 15 latent dimensions for ClearSMILES (orange), RDKit Default canoncial SMILES, canonical kekule SMILES

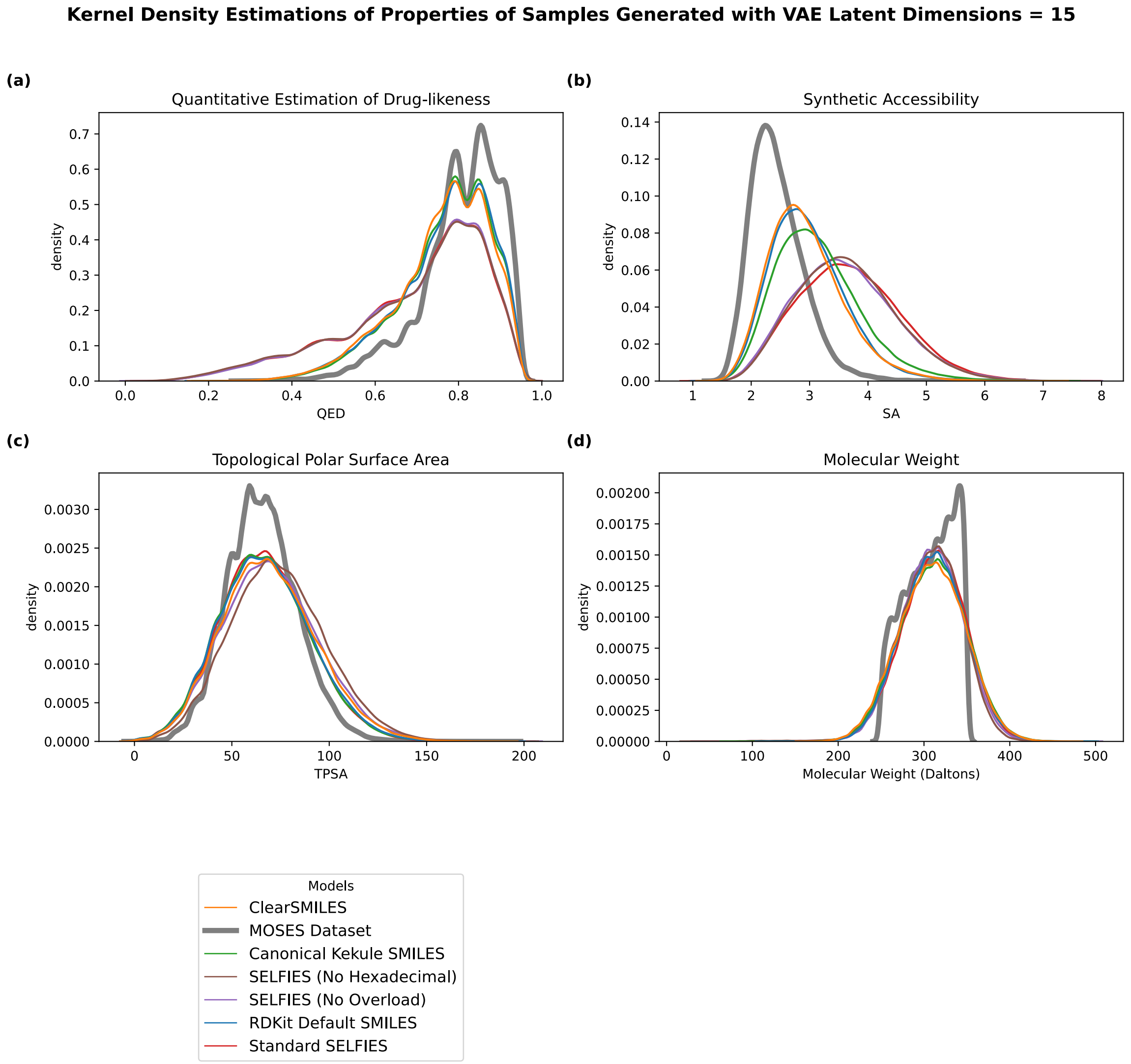


Figure 19: Assessment of various fidelity metrics for all viable samples generated by all VAE models with 15 latent dimensions: (a) QED, (b) SA, (c) TPSA, (d) molecular weight

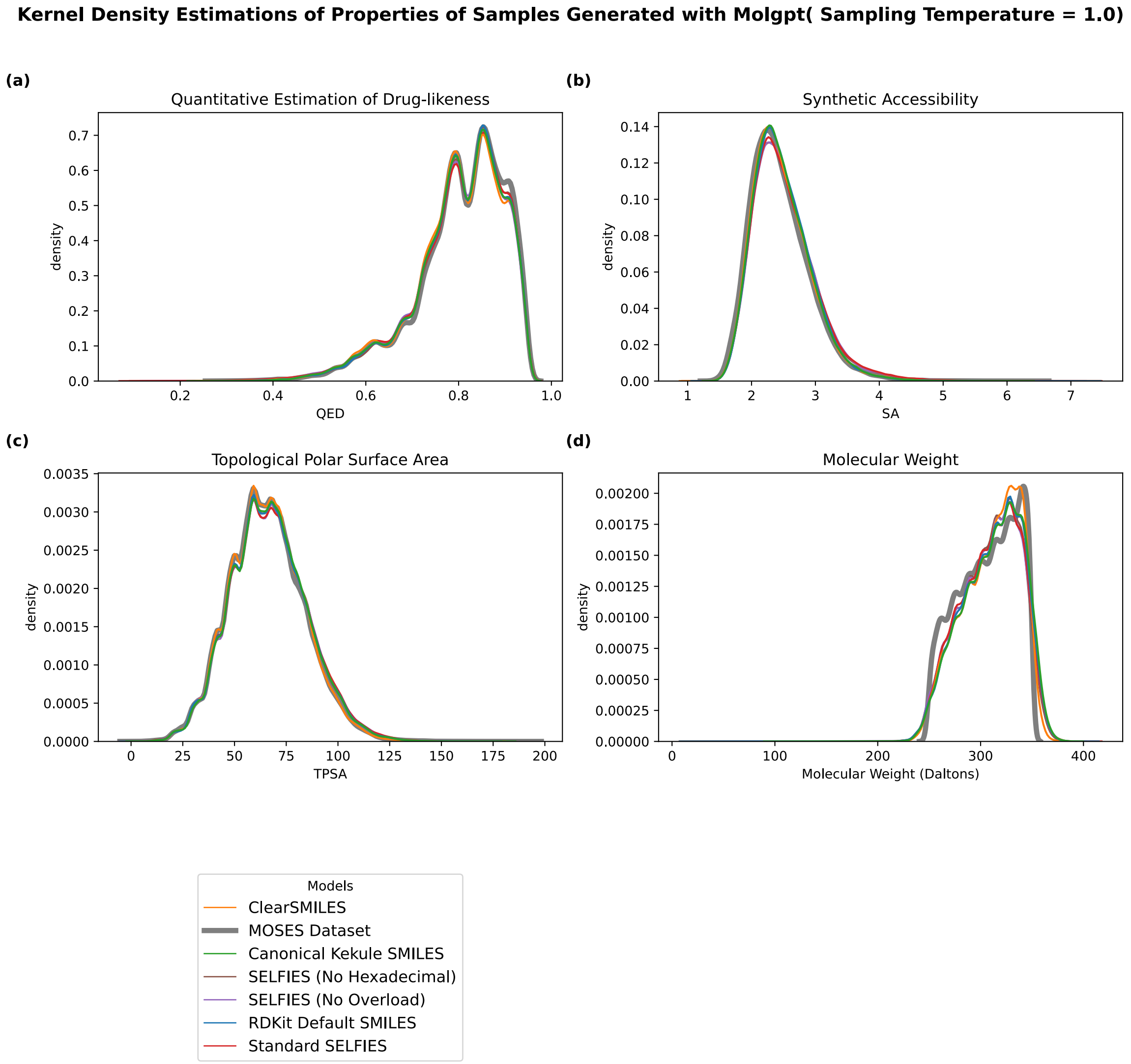


Figure 20: Assessment of various fidelity metrics for all viable samples generated by MolGPTs models with sampling temperature of 1.0: (a) QED, (b) SA, (c) TPSA, (d) molecular weight
